# Structural Insights on Ionizable Dlin-MC3-DMA Lipids in DOPC Layers by Combining Accurate Atomistic Force Fields, Molecular Dynamics Simulations and Neutron Reflectivity

**DOI:** 10.1101/2023.02.28.529897

**Authors:** Mohd Ibrahim, Jennifer Gilbert, Marcel Heinz, Tommy Nylander, Nadine Schwierz

**Affiliations:** Department of Theoretical Biophysics, Max Planck Institute of Biophysics, Max-von-Laue-Straße 3, 60438 Frankfurt am Main, Germany; Institute of Physics, University of Augsburg, 86159 Augsburg, Germany; Physical Chemistry, Department of Chemistry Lund University P.O Box 124, SE-22100 Lund, Sweden

## Abstract

Ionizable lipids such as the promising Dlin-MC3-DMA (MC3) are essential for the successful design of lipid nanoparticles (LNPs) as drug delivery agents. Combining molecular dynamics simulations with experimental data such as neutron reflectivity experiments and other scattering techniques is essential to provide insights into the internal structure of LNPs, which is not fully understood to date. However, the accuracy of the simulations relies on the choice of force field parameters and high-quality experimental data is indispensable to verify the parametrization. For MC3, different parameterizations in combination with the CHARMM and the Slipids force field have recently emerged. Here, we complement the existing efforts by providing parameters for cationic and neutral MC3 compatible with the AMBER Lipid17 force field. Subsequently, we carefully assess the accuracy of the different force fields by providing a direct comparison to neutron reflectivity experiments of mixed lipid bilayers consisting of MC3 and DOPC at different pH. At low pH (cationic MC3) and at high pH (neutral MC3) the newly developed MC3 parameters in combination with AMBER Lipid17 for DOPC give good agreement with the experiments. Overall, the agreement is similar compared to the Park-Im parameters for MC3 in combination with the CHARMM36 force field for DOPC. The Ermilova-Swenson MC3 parameters in combination with the Slipids force field underestimate the bilayer thickness. While the distribution of cationic MC3 is very similar, the different force fields for neutral MC3 reveal distinct differences ranging from strong accumulation in the membrane center (current MC3/AMBER Lipid17 DOPC), over mild accumulation (Park-Im MC3/CHARMM36 DOPC) to surface accumulation (Ermilova-Swenson MC3/Slipids DOPC). These pronounced differences highlight the importance of accurate force field parameters and their experimental validation.

---

Ionizable lipids are the key component of clinically translatable lipid nanoparticles (LNPs). With the incredible growth in RNA therapeutics, LNPs have become an indispensable tool to deliver RNA to target cells thereby providing promising perspectives to combat life-threatening diseases such as Amyloidosis or COVID-19^1–3^. In addition to neutral helper lipids, cholesterol, and PEG lipids, ionizable lipids are the key components since they attach to and protect the RNA cargo and facilitate cytosolic transport ^4^. Here, the positive charge of the lipids at acidic pH condenses the RNA while the lack of charge at physiological pH minimizes toxicity. To date, several ionizable lipids have been successfully developed ^5^. MC3 was identified as one of the most promising ionizable lipids due to its high transfection efficiency ^2,4,6^ and is used in the first FDA-approved drug for the treatment of amyloidosis ^3^. Despite their importance, LNPs are still limited by low transfection or release efficiencies. Here, a detailed understanding of the distribution of the ionizable lipids within the LNP, their interactions with the RNA cargo and their role during endosomal disruption is required for further progress.

Molecular dynamics simulations are particularly suited to resolve such interactions and to provide atomic insights into the structure. However, the accuracy of the simulations depends on the empirical force fields which describe the interactions of the atoms in the system. Out of the large variety, three force field families are most commonly used in biomolecular simulation of lipids: (i) The CHARMM36 lipid force field ^7^, (ii) the Slipids ^8^ and (iii) the AMBER Lipid14^9^ and Lipid17^10^ force fields. The force fields are derived using different parametrization strategies using higher level quantum mechanical calculations and validated by experimental results such as the area per lipid, volume per lipid, isothermal compressibility, NMR order parameters, scattering form factors, and lipid lateral diffusion coefficient. For lipid-only simulations, CHARMM36 and Lipid14/17 are considered to give the closest agreement with experiments ^11^. In addition, the force fields are considered to be compatible within their families since they are derived based on the same parameterization strategy. For instance, the Lipid17 force field can be combined easily with the AMBER force fields for proteins, small molecules, nucleic acids, and carbohydrates, which greatly enhances its use in the field of biomolecular simulations. Despite the consistent parameterization within one family, we may note that it is still recommended to carefully check whether such combinations lead to physically meaningful results for the system under consideration ^12,13^.

In 2020, Ermilova et al. derived parameters for MC3 in combination with the Slipids force field ^14^. Unfortunately, the authors did not parametrize the protonated state of the MC3 molecule. Since MC3 has an apparent pKa of about 6.44^6^the protonated state plays an essential role: It ensures efficient encapsulation of the nucleic acids at low pH and a high charge at endosomal pH enabling the drug release from the LNP. At physiological pH, the MC3 lipids are neutral enabling circulation of largely uncharged LNPs. For the modeling it is therefore beneficial to parametrize both states simultaneously.

More recently, Park et al. derived parameters for neutral and protonated MC3 in combination with the CHARMM36 lipid force field ^15^. However, to assess the accuracy of the different force fields, a direct comparison with experimental data is required.

The aim of our study is to provide accurate parameters to correctly model complex systems such as LNPs. However, to date it is impossible to provide a reliable quantitative comparison between all-atom MD simulations and scattering experiments for such large, multi-component systems. In order to validate atomistic models, we therefore use lipid bilayers consisting of mixtures of MC3 and DOPC as valuable model systems which allows us to directly compare experiments and simulations.

X-ray and neutron scattering are one of the most reliable experimental techniques to characterize such systems ^16–18^. Neutron reflectivity is particularly suited to characterize the interfaces on the nanometer length scale ^19,20^. Neutrons are beneficial due to their non-destructive nature, large penetration depth, and sensitivity to the nuclear scattering length which enables specific parts of the system to be highlighted by replacing hydrogen with its heavy isotope deuterium. The neutron scattering length density (SLD) contains information on the molecular composition of the system perpendicular to the interface ^21–23^. However, from the experimentally measured reflectivity profiles, an iterative modeling approach is typically required to obtain the SLD profiles. It is therefore particularly appealing to combine neutron reflectivity experiments and MD simulations to make robust and model-free predictions. The MD simulations allow us to calculate the SLD and hence the reflectivity profile without further assumptions and to compare directly to the reflectivity experiments. Similar strategies have been used in previous works to characterize the structure of biological membranes and membrane-protein systems ^24–26^.

The aim of the current work is to provide insights into the structure of lipid bilayers consisting of MC3 and DOPC at different pH by combining molecular dynamics simulations and neutron reflectometry experiments. To that end, we first derive force field parameters for protonated and neutral MC3 in combination with the AMBER Lipid17 DOPC force field. We perform simulations using the currently derived as well as existing MC3 force fields. The direct comparison of the simulations with neutron reflectivity experiments at two different pH values and three different solvent contrasts allows us to carefully assess the predictions of different force fields.

## 1. Methods

### 1.1 Computational methods

Parameterization of MC3: We first derive force field parameters for protonated and neutral MC3 in combination with the AMBER Lipid17 DOPC force field (see section 2.1 in the Supporting Information). The force field parameters are available at https://github.com/bio-phys/ForceFieldsMC3. MD simulations: We simulated DOPC/MC3 mixtures at two different pH values with the mole fraction of MC3 varied from 5%-15%. Based on the pKa value of 6.44 for MC3 obtained by in situ fluorescence titration of LNPs ^6^ and the Henderson-Hasselbalch equation, the majority of MC3 molecules is charged at pH 6 (73 %) while most of them are uncharged at pH 7 (78 %). Note that the pKa of MC3 in lipid DOPC layers might deviate from the one in LNPs due to the differences in composition and structure. However, small changes of about 20% in the degree of protonation did not significantly change the calculated reflectivity profiles (Figure S8). We have therefore chosen to compare the systems that are fully charged at pH 6 and uncharged at pH 7, which represent the limiting cases.

In total, we used 5 different force field combinations: (i) Park-Im MC3/MC3H and CHARMM36 DOPC ^15^, (ii) Ermilova-Swenson MC3^14^ and Slipids DOPC ^8^, (ii) current MC3/MC3H and AMBER Lipid17 DOPC ^10^. The CHARMM-GUI web-server ^27^ was used to generate the initial configuration in the CHARMM-based simulations and the MemGen web-server ^28^ otherwise. The bilayer contains 100 lipids per leaflet for the setups with 15% MC3 and 200 lipids per leaflet for lower MC3 mole fractions. Further details on the simulation setups are given in Table S1. For the CHARMM-based systems, the mTIP3P ^29^ water model was used and the TIP3P ^30^ water model otherwise. A physiological salt concentration of 150 mM NaCl was used and extra ions were added to neutralize the systems with MC3H. For the ions, the MamatklovSchwierz force field parameters optimized in combination with TIP3P water were used ^31^. Further details on the MD simulations are given in the supporting information (section 2.2). The simulations were performed for 600 ns. To test the convergence of the simulations, we performed a block analysis ^32^ to calculate the variance of the average area per lipid indicating converged results after about 75 ns (Figure S9).

To compare simulations and experiments, we calculated the reflectivity profiles. The last 300 ns was used to calculate the SLD profiles. To provide a direct quantitative comparison, the silicon substrate not present in the simulations was included in a three-step modelling procedure. First, the silicon substrate is modeled by fitting the experimental data of the substrate in solution without the bilayer to a three-slab model (see section 2.3.1 and Figure S3 in the supporting information). The roughness was allowed to vary during the fitting and the optimum values were found to be between 1-2 Å for each interface. Secondly, the position of the substrate/bilayer interface is determined from the SLD profile from the simulations (see section 2.3.2 and Figure S4 in the supporting information). We assume that the bilayer and the substrate are in contact similar as in previous work ^24,25^. Thirdly, we fitted the water fraction and water patches by introducing two fit parameters. *α* corresponds to the amount of water in the bilayer leaflet closer to the substrate. The second parameter *γ* corresponds to the fraction of water patches on the substrate when it is not perfectly covered by the bilayer (see Figure S5 A-B in the supporting information for an illustration of the effect of *α* and *γ* on the SLD profile). Finally, the two parameters are optimized using a grid search to obtain the global minimum of *χ*^2^ for all three contrasts (see supporting information, section 2.3.4). Abele’s matrix formalism as implemented in the Refnx ^33^ python package was used to obtain reflectivity curves from the SLD profiles.

### 1.2 Experimental methods

#### 1.2.1 Materials

1,2-dioleoyl-sn-glycero-3-phosphocholine (DOPC, powder, >99%) was purchased from Avanti Polar Lipids (Birmingham, AL, USA) and (6Z,9Z,28Z,31Z)-heptatriacont-6,9,28,31-tetraene19-yl 4-(dimethylamino)butanoate (DLin-MC3-DMA or MC3, liquid oil, >98%) was purchased from Biorbyt (Cambridge, UK). Chloroform (>99.8%), buffer salts (NaCl, KH_2_PO_4_, Na_2_HPO_4_, NaH_2_PO_4_.H_2_O, all with purity >99.0%) and D_2_O (>99.9% D atom) were purchased from Sigma Aldrich. Milli-Q purified water (18 MΩ cm) was used for all experiments. 50 mM sodium phosphate buffers (pH6 and pH7) were prepared in D_2_O, H_2_O and a mix of 38:62 D_2_O:H_2_O (CMSi, contrast matched to Si) by mixing the corresponding 50mM Na_2_HPO_4_ and 50mM NaH_2_PO_4_.H_2_O solutions while monitoring the pH to the target pH.

#### 1.2.2 Preparation of lipid vesicles

Lipid vesicles were prepared using a modified version of the protocol used by Dabkowska et al. ^34^. In summary, lipid stock solutions were prepared in chloroform and mixed to the required molar ratio. The chloroform was then evaporated under N_2_ flow and the lipids films were desiccated for 14hrs to remove any remaining solvent. For in house ellipsometry measurements, the lipid film was then hydrated in phosphate buffered saline (1x PBS, pH 7.2*±*0.05, 155 mM NaCl, 2.97 mM Na_2_HPO_4_, 1.06 mM KH_2_PO_4_), vortexed until all lipid film had been removed from the surface of the vial and left for 15 mins. The dispersion, with a lipid concentration of 0.5 mg/mL was sonicated in an ice water bath using a tip sonicator (Vibra-Cell VCX 130, Sonics & Materials Inc., Newton, CT, USA) with the following settings: 15 mins sonication time, 10s sonication followed by a 10s cooling period, 50% amplitude. The sizes of the vesicles produced were measured using dynamic light scattering (DLS) and are presented in Table S3.

The neutron reflectometry (NR) experiment was performed remotely, therefore the lipid films were prepared 1 week in advance, shipped in dry ice and stored at -20 °C. For the NR experiment, sample preparation was as described above, except the lipid films were hydrated to 2 mg/mL before sonication using a Fisherbrand Model 50 Sonic Dismembrator (FisherScientific, Waltham, MA, USA) and diluted to 0.5 mg/mL in 1x PBS before injection into the measurement cell.

#### 1.2.3 Neutron Reflectrometry

Neutron reflectometry measurements were performed using polished silicon substrates (Sil’Tronix, Archamps, France) with the dimensions 50mm x 80mm x 15mm. The substrates were cleaned using the RCA method, thoroughly rinsed with MilliQ and dried using N_2_, then immediately introduced to the custom reflectometry cells. Specular neutron reflectometry measurements were performed on systems containing 15% MC3 and 85% DOPC at two different pH conditions, pH = 6 and pH = 7. The measurements were performed on the POLREF reflectometer at the ISIS Neutron and Muon Source (STFC Rutherford Appleton Laboratory, Didcot, UK) over q range 0.009 - 0.27 Å^*−*1^ at 25 °C using a custom built cell as described in detail elsewhere. The measurements proceeded as follows: (i) characterisation of bare surface in D_2_O and H_2_O, (ii) manual injection of vesicles (incubation for 45 mins 60 mins), (iii) manual rinsing with MilliQ, (iv) characterisation of lipid layer in 50mM sodium phosphate buffer in 100% D_2_O buffer, 100% H_2_O buffer and contrast matched silicon (CMSi) buffer (38 % D_2_O). The raw data was reduced using Mantid Workbench ^35^.

### 1.2.4 Ellipsometry

Ellipsometry measurements were performed using a UVISEL spectroscopic ellipsometer (HORIBA Jobin Yvon/HORIBA Scientific, Middlesex, UK). The changes in polarisation in terms of the amplitude ratio, Ψ, and phase difference, Δ, were measured using an angle of incidence of 70°, a modulator angle of 0°, and an analyser angle of 45°. The beam diameter used was 1.2 mm, resulting in a spot size of 2.1 mm at the interface. All experiments were performed at 25°C using a wavelength range of 190 - 824 nm. The thickness of the lipid layer was measured by in situ spectroscopic ellipsometry using a custom built cuvette, which allows exchange of the solution without emptying the cuvette. Details on the set-up are given in Humphreys et al. ^36^. Initially the dry silicon substrate was characterised, then again after hydration with 1x PBS. 1 mL of vesicle sample was manually injected into the cell, left to incubate and rinsed with Milli Q, then 50 mM sodium phosphate buffer using a peristaltic pump with a flow rate of 2.25 mL/min. At each stage the equilibrated sample was characterised after the measured Ψ and Δ values stabilised.

Data analysis to calculate the layer thickness was performed using the refellips software ^37^, which is analogous to the refnx ^33^ package for the fitting of neutron and x-ray scattering data. The theoretical model used optical constant values derived from literature for silicon, silica, air and water ^38–42^. The Cauchy equation can be used to account for the wavelength dependent nature of the refractive index:

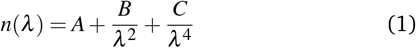

where n(*λ*) is the wavelength dependent refractive index and A, B and C are coefficients characteristic of the material. Here the values for a hydrated lipid in the DeltaPsi2 software (HORIBA Jobin Yvon/HORIBA Scientific, Middlesex, UK) were used: A = 1.45, B = 0.01, C = 0. A multilayer slab model was used to model the lipid data, composed of silicon (backing) a hydrated silica layer (slab 1), an effective medium approximation (EMA) layer for the hydrated lipid layer (slab 2) and water (for all other measurements). The thickness of the silica layer was determined and consequently fixed by fitting the bare silica data. The lipid layer was modelled as a single homogeneous slab with an associated roughness (fixed = 0 Å), volume fraction of solvent (fixed = average volume fraction of solvent over layer from simulation using current MC3/AMBER Lipid17 DOPC) and thickness (varied). It should be noted that when the roughness was varied, there was no significant change in the fit result. The data was fit in the wavelength range 320 - 650 nm using both least squares and differential evolution, which produced the same thickness value within error.

The adsorbed mass of the layer was additionally calculated from the fitted thickness and volume fraction of solvent using the following equation:

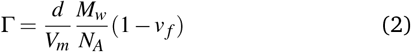

where Γ is adsorbed mass (mg m^*−*3^), V_*m*_ is molecular volume, M_*w*_ is molecular weight, N_*A*_ is Avogadro’s number, v _*f*_ is the volume fraction of solvent in the layer. Here, V_*m*_ is the weighted average molecular volume of MC3 and DOPC, M_*w*_ is the weighted average molecular weight of MC3 and DOPC, and v _*f*_ is the average volume fraction of solvent over layer from simulation using current MC3/AMBER Lipid17 DOPC.

## 2 Results and discussion

### 2.1 Distribution of neutral and cationic MC3 in lipid bilayers

The simulations of bilayers consisting of DOPC and neutral MC3 reveal that different force fields have a significant influence on the distribution of MC3 in the bilayer (Figure 1A). With the force field parameters derived in our current work, the neutral MC3 molecules segregate partially from the DOPC lipids and accumulate in the center. The accumulation leads to an increase in bilayer thickness from 36.8 Å for pure DOPC to 41.5 Å for 15 mol % MC3 (see Tables S4, S5).

**Fig. 1.**
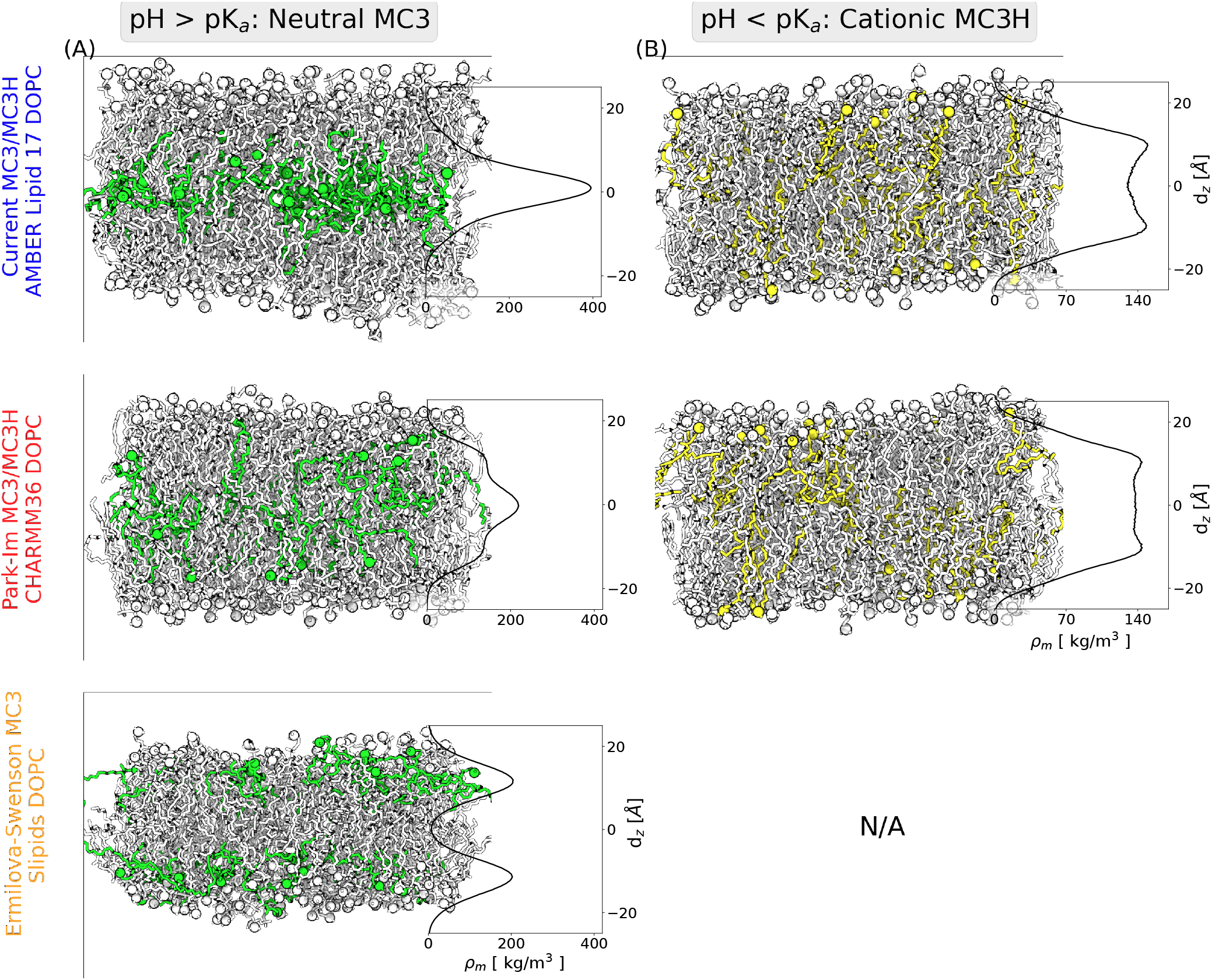
Simulation snapshots and mass density profiles of 15% MC3 in a DOPC bilayer for the different force fields at high pH where MC3 is neutral (A) and at low pH where MC3 is cationic (B). The spheres represent the nitrogen atoms in the head groups. Neutral MC3 is shown in green, cationic MC3 in yellow, and DOPC in white. The simulations were performed with five different force fields: (i) Park-Im MC3/MC3H and CHARMM36 DOPC ^7^, (ii) Ermilova-Swenson MC3 and Slipids DOPC ^8^, (ii) current MC3/MC3H and AMBER Lipid17 DOPC ^10^.

With the Park-Im MC3 and CHARMM36 DOPC parameters, the accumulation is less pronounced and the and the bilayer thickness increases only slightly (Table S4). This behavior can be rationalized by the fact that in the Park-Im MC3 parameterization, the nitrogen has a larger partial charge compared to the current parameterization. The electrostatic interactions with the carbonyl group of DOPC are therefore stronger as reflected in the radial distribution function (Figure S12).

For the Ermilova-Swenson MC3 in combination with the Slipids DOPC parameters, the neutral MC3 lipids accumulate at the lipid/water interface. The partitioning of the hydrophobic tails near the hydrophilic head group region is surprising and possibly arises from the differences in the Lennard-Jones or other nonbonded force field parameters. Overall the accumulation of MC3 in the interfacial region results in a significant decrease of the bilayer thickness compared to a pure DOPC bilayer from 36.7 Å for pure DOPC to 33.2 Å for 15 mol % MC3 (see Table S4). Given the variety of distributions and bilayer thicknesses obtained from different force field parameters the immediate question arises which parameters should be chosen to obtain reliable results.

Initial insight is obtained from the behavior of DLin-KC2-DMA (KC2) which is similar to MC3. Recent simulations and cryo-TEM and SAXS experiments ^43^ show that neutral KC2 has a high tendency to segregate from POPC. In addition, experiments on LNPs containing MC3 or KC2 show a high electron density core ^44,45^ which is attributed to neutral MC3/KC2 forming an oil droplet in the LNP core. The simulations with the current force field for MC3 or the Park-Im MC3 parameters agree with such behavior. By contrast, the accumulation of MC3 at the bilayer-water interface for the Ermilova-Swenson force field does not seem to explain such results.

The experimental reflectivity measurements, obtained at two different pH values, offer further insights into the structure of the lipid layer. The presented experimental reflectivity data are extracted from an ongoing study focused on the interaction between nucleic acid a lipid layers with MC3^46^. At 100% D_2_O, which provides the largest contrast between solvent and the hydrocarbon tails, a clear shift of the characteristic minimum to lower *q* values is observed (Figure S13) and reflects an increase of the bilayer thickness for neutral MC3 compared to cationic MC3. The increase of the bilayer thickness is further supported by the shift of the minimum in the reflectivity profiles to lower q upon increasing the MC3 mol fraction from 5 % to 15 % (Figure S14). Note that these tendencies are obscure in the ellipsometry data due to the large uncertainties in the measurements. (Table 1)

**Table 1.**
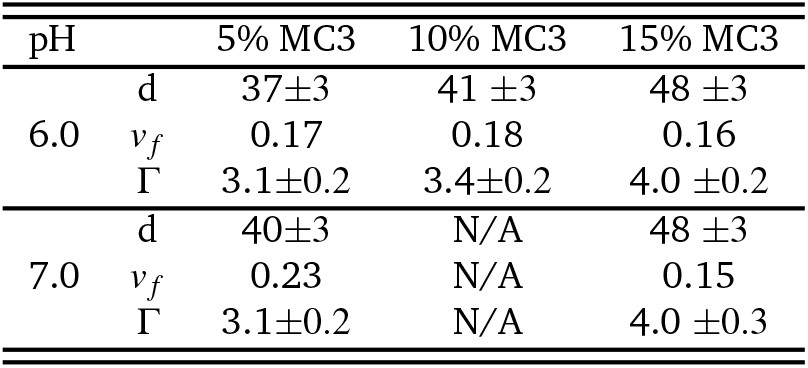
Ellipsometry measurements: Thickness, d (Å), and adsorbed mass, Γ (mg m^*−*3^), of MC3/DOPC layers from ellipsometry measurements in 50mM sodium phosphate buffer at pH 6 or 7. The solvent volume fractions, *v* _*f*_, used to calculate these values were extracted from simulations using the current MC3/MC3H and Lipid 17 force fields.The error given is the error from the refellips fitting of the single slab model of the lipid bilayer to the data based on *χ*^2^ mininimization of deviation between model and the experimental data as described in ^37^.

| pH |  | 5% MC3 | 10% MC3 | 15% MC3 |
| --- | --- | --- | --- | --- |
| 6.0 | d | 37±3 | 41 ±3 | 48 ±3 |
| | $v_f$ | 0.17 | 0.18 | 0.16 |
| | $\Gamma$ | 3.1±0.2 | 3.4±0.2 | 4.0 ±0.2 |
| 7.0 | d | 40±3 | N/A | 48 ±3 |
| | $v_f$ | 0.23 | N/A | 0.15 |
| | $\Gamma$ | 3.1±0.2 | N/A | 4.0 ±0.3 |

For the bilayer simulations, consisting of cationic MC3H and DOPC, the differences between the two force fields are less pronounced (Figure1). In both cases, the charged MC3H head groups remain at the lipid/water interface, as expected, and the thickness does not change significantly (Table S4). Overall, the Park-Im MC3H parameters have a slightly higher tendency toward the bilayer center as evident from mass density profiles of MC3H (Figure 1B). Overall, the observations for MC3H are similar to the results for protonated KC2H ^43^.

### 2.2 Direct comparison of simulations and neutron reflectivity experiments

In the following, we compare the reflectivity profiles obtained from all-atom MD simulations with the experimental measurements in different solvent contrasts. However, to provide a direct quantitative comparison between experimental and simulation results, the influence of the solid support, which is not included in the simulations, has to be considered. This is achieved in a three-step procedure: The silicon substrate is modeled by fitting the experimental data of the substrate in solution to a three-slab model (Figure S3). Subsequently, the position of the solid/bilayer interface is determined from the simulations (Figure S4). Finally, we follow the procedure of previous work ^24,25^ and introduce two physically motivated fit parameters. The first parameter, *α*, accounts for the content of water in the leaflet next to the substrate. The second parameter, *γ*, accounts for possible water patches due to imperfect substrate coverage. Note that the phospholipid bilayer surface coverage on silicon substrate can be as low as 70% ^47–49^.

In summary, the three-step procedure introduces a minimum amount of parameters. The parameters of the three-slab model are obtained from additional measurements containing only the substrate. Two additional fit parameters are necessary to indirectly take into account the lipid bilayer coverage on the substrate and hydration level in the leaflet next to the substrate. The procedure allows us to assess the accuracy of the different force fields for MC3 and DOPC through the deviations between experimental and simulation results as discussed in the following.

#### 2.2.1 Cationic MC3H-DOPC bilayers: Comparison of reflectivity profiles from simulations and experiments

Figure 2 shows the neutron scattering length density (SLD) and the reflectivity profiles for mixed MC3H/DOPC bilayers from experiments and simulations at pH 6 at three different solvent contrasts. In the simulations, we assume that all MC3 molecules are protonated (see Methods section 1.1). Both force fields give good agreement with the experiments at all solvent contrasts as evident from visual inspection (Figure 2D-F) and from low *χ*^2^ values (Figure S7).

**Fig. 2.**
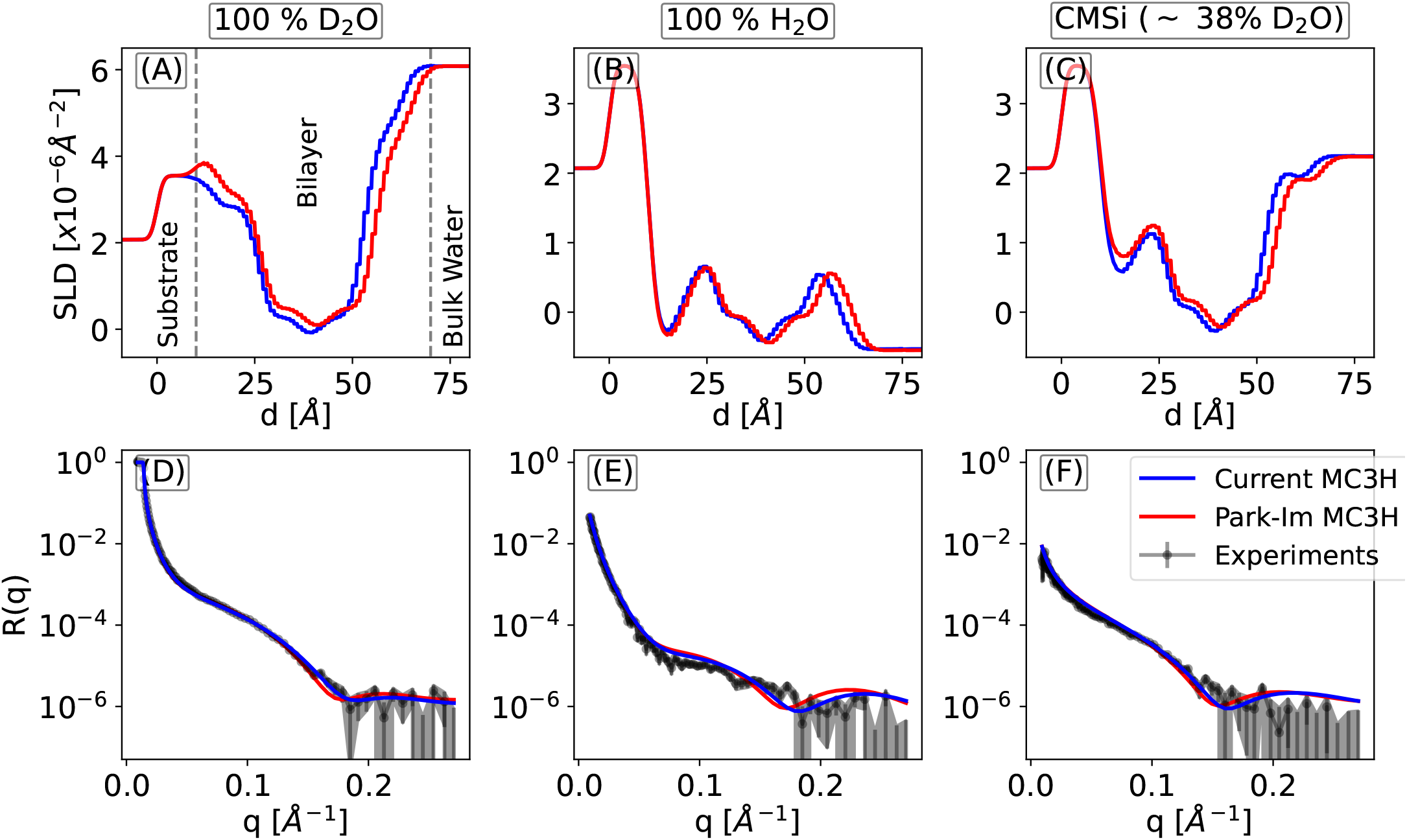
Direct comparison of simulations and neutron reflectivity experiments for 15% cationic MC3 in a DOPC bilayer. Neutron scattering length density profiles from the simulations with two different force fields at three different deuteration levels (top) and reflectivity profiles from simulations and experiments (bottom). The contrast corresponds to 100% D_2_O (A), 100 % H_2_O (B) and contrast matched silicon at *∼* 38% D_2_O. The dashed lines in (A) indicate the regions of substrate, bilayer, and water. Simulation snapshots for the two force fields are shown in Figure 1 B. The experiments were performed at pH 6 and the error bars are calculated from error propagation of the square root of the number of counts per bin on the detector during the data reduction. The larger error bars at high q are expected as the effect of the signal from the reflection is lower and approaches the level of the background scattering.

The SLD profiles also reflect the slightly different distributions of cationic MC3H in the bilayer DOPC (Figure1B). However, such subtle differences are lost in the resulting reflectivity profiles due to the convoluted contributions of MC3, DOPC, water, and substrate.

The current force field parameters for MC3H in combination with AMBER Lipid17 also give good agreement at lower MC3H fractions for all solvent contrast (Figure S15 for 5% MC3H mol fraction and Figure S16 for 10% MC3H mol fraction).

The substrate parameters obtained for the two different force fields are consistent and we obtain similar values: *α* = 0.53, *γ* = 5.0% (current MC3H and AMBER DOPC) and *α* = 0.62, *γ* = 8.0% (Park-Im MC3H and CHARMM36 DOPC). In addition, the fraction of uncovered substrate area obtained for different MC3H mole fractions yield *γ* = 5.0%-11.6%, which is well within the range of values reported from experiments ^47–49^ and simulations ^24,25^. The consistent values further substantiate the robustness of the substrate modeling.

#### 2.2.2 Neutral MC3-DOPC bilayers: Comparison of reflectivity profiles from simulations and experiments

Figure 3 shows the SLD and the reflectivity profiles for mixed MC3/DOPC bilayers from experiments and simulations at pH 7 and three different deuteration levels. In the simulations, we assume that all MC3 molecules are neutral (see Methods section 1.1). The results with the Park-Im and the current parameters are similar and agree well with the experimental results (Figure 3D-F). In particular, the position of the characteristic minimum, clearly visible at 100% D_2_O, is well captured. However, the results with the Ermilova-Swenson MC3 parameters deviate both in the SLD and the reflectivity profile. This is expected as their model predicts a different distribution of MC3 in the lipid bilayer with MC3 accumulated at the lipid/water interface (Figure 1A). The comparison with experiments reveals that the characteristic minimum is shifted to higher *q* values indicating that the combination of Ermilova-Swenson MC3 and Slipids DOPC parameters underestimates the bilayer thickness. Again, this is likely due to the accumulation of neutral MC3 at the lipid/water interface. In contrast, the other two models, in which neutral MC3 accumulates in the middle of the bilayer, are better suited to describe the experimental data.

**Fig. 3.**
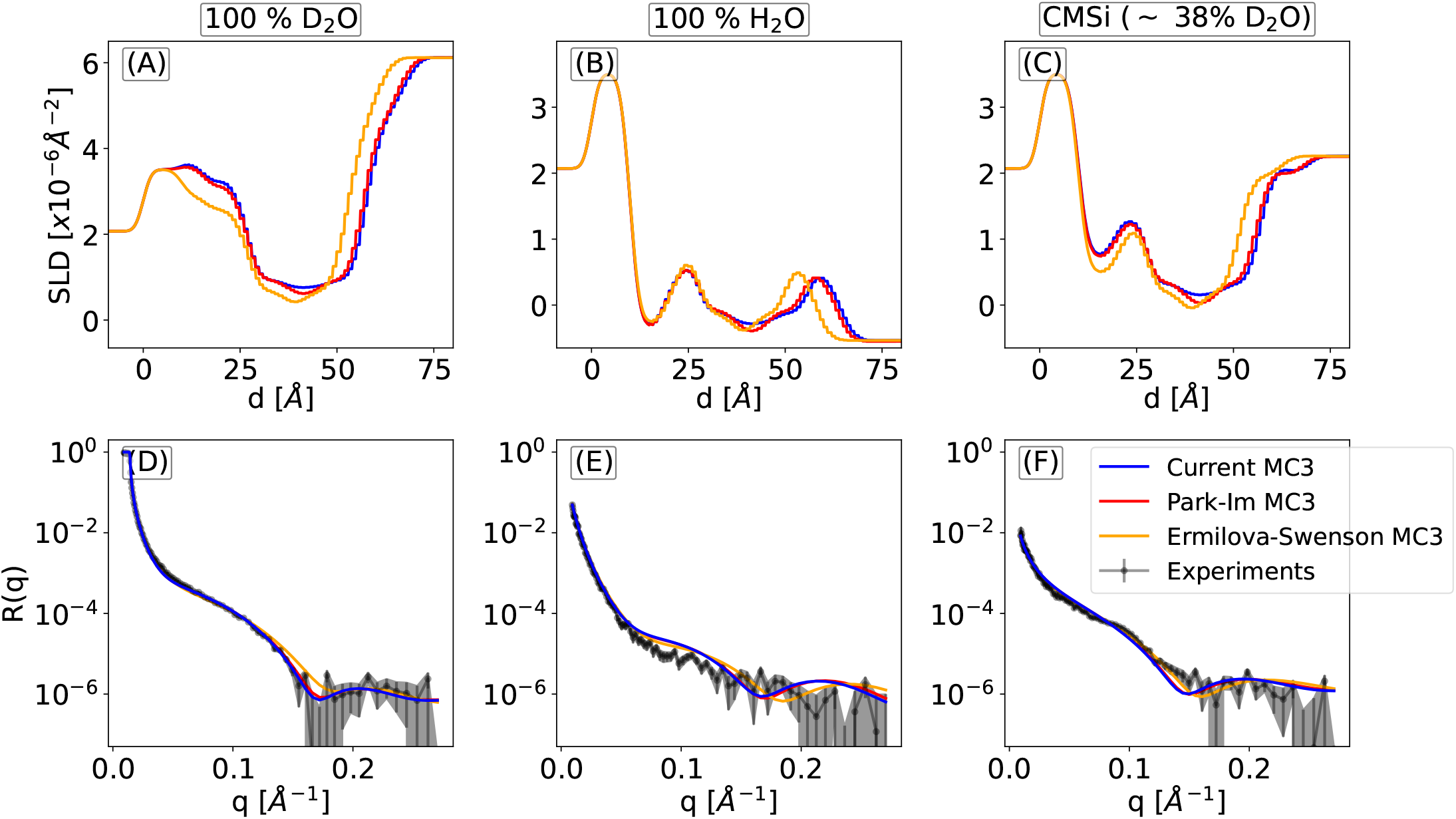
Direct comparison of simulations and neutron reflectivity experiments for 15% neutral MC3 in a DOPC bilayer. Neutron scattering length density profiles from the simulations with three different force fields at three different deuteration levels (top) and reflectivity profiles from simulations and experiments (bottom). The contrast corresponds to 100% D_2_O (A), 100 % H_2_O (B) and contrast matched silicon at *∼* 38% D_2_O. The experiments were performed at pH 7. The corresponding simulation snapshots are shown in Figure 1 A.

As before, the substrate parameters are consistent. For the three force fields we obtain similar values: *α* = 0.52, *γ* = 15.6% (current MC3 and AMBER DOPC), *α* = 0.51, *γ* = 15.1% (ParkIm MC3H and CHARMM36 DOPC) and *α* = 0.40, *γ* = 12.1% (Ermilova-Swenson MC3 and Slipids). In addition, the fraction of uncovered substrate area obtained from the independent fits at the different pH conditions yields *γ* = 12%-15.6%. The uncovered substrate area is slightly higher for neutral MC3 compared to cationic MC3H.

With the current force field parameters for neutral MC3 in combination with AMBER Lipid17 DOPC, we also obtain good agreement with the experiments at lower MC3 fraction in all solvent contrasts (Figure S17 for 5% MC3H mol fraction).

Overall, the agreement between experiments and simulations with the current force fields is consistent and good both for cationic and neutral MC3 in a DOPC bilayer. Still, small deviations in the reflectivity profiles are observed (Figure 2D-F and Figure 3D-F). Possible sources for the deviations are (i) explicit substrate effects not covered in the current modeling procedure, (ii) different degrees of protonation in experiments and simulations at pH 7, and (iii) inaccuracies of the atomistic force fields.

Regarding the substrate effect, our current modeling does not include any structural changes of a free bilayer compared to a solid-supported one. The presence of a solid support may influence both the interfacial water density and the bilayer SLD ^25^. However, considering the good agreement at low pH, the effect of such structural changes on the measured reflectivity profiles is expected to be small. Explicit modeling of the substrate would introduce a large number of additional force field parameters with unknown accuracy.

To test the influence of a different degree of protonation at pH 6 or pH 7, we performed simulations with varying degrees of protonation but did not observe a significant change in the results (Figure S8). It is therefore most likely that the small deviations are caused by shortcomings of the atomistic force fields of water, DOPC, and/or MC3. Here, the question arises which force field component has the largest effect and how it can be improved further.

In the current work, we used the TIP3P (AMBER) or mTIP3P (CHARMM) water model ^29,30^ due to their compatibility with the respective lipid force fields. However, a large variety of water models exists, which reproduce the experimental results for the structural and physical properties of water much better ^50^. Superior water models might improve the agreement but also modify the lipid-water interactions.

Surprisingly, the force field parameters for neutral MC3 from the current approach and the Park-Im MC3/CHARMM DOPC parameters yield very similar reflectivity profiles (Figure 3D-F) even though the distribution of the ionizable lipid in the DOPC bilayer is rather different (Figure 1A). Hence, strong, or mild accumulation of neutral MC3 in the bilayer reproduces the experimental data equally well, possibly due to a compensation of the individual contributions of MC3 and DOPC. Even though the experimental neutron reflectometry profiles indicate an increase of the bilayer thickness (Figure S14) and support the hypothesis of an accumulation of MC3 as predicted by the current parameters, additional experiments are required to determine the correct magnitude of segregation. For example, the form factors calculated for the three distributions of MC3 shown in Figure 1A reveals pronounced differences. Small angle X-ray scattering (SAXS) experiments could therefore resolve this problem (Figure S18) and serve as a starting point for further optimization. A promising starting point for such optimization are the angles and dihedrals currently not present in the Lipid17 force field. Moreover, we chose the HF/6-31G* basis set to derive the partial charges as in AMBER Lipid14. This choice is validated by comparing to experimental reflectometry data. However, a larger basis set or implicit solvent model might lead to slightly different partial charges and possibly improved agreement with experimental data.

## 3 Summary

Ionizable lipids play a crucial role in the development of lipid nanoparticles for drug delivery. Molecular dynamics simulations are ideal to resolve the microscopic structures but rely on accurate atomistic force fields. Moreover, to correctly describe the interactions in such multicomponent systems, the force fields for neutral and ionizable lipids must be compatible.

In this current work, we developed force field parameters for the ionizable lipid Dlin-MC3-DMA for biomolecular simulations in combination with the AMBER force field. To validate existing and newly developed force field parameters for cationic and neutral MC3, we performed neutron reflectivity experiments for bilayers consisting of DOPC and MC3 at three different solvent contrasts, two different pH values, and different fractions of MC3. Our results show that combining MD simulations and neutron reflectivity experiments is particularly suited to assess the accuracy of the force fields. At low pH (cationic MC3), the current MC3H parameters in combination with AMBER Lipid17 for DOPC and the Park-Im MC3H parameters in combination with CHARMM36 for DOPC yield similar distributions and good agreement with the experimental neutron reflectivity profiles. At high pH (neutral MC3), the three different force fields lead to clearly different distributions of MC3 in the DOPC bilayer: The Ermilova-Swenson MC3 parameters in combination with the Slipids DOPC predict an accumulation of MC3 at the lipid/water interface. The Park-Im MC3 parameters in combination with CHARMM36 for DOPC yield a mild accumulation in the bilayer center. The current MC3 parameters in combination with AMBER Lipid17 for DOPC yield a stronger accumulation in the bilayer center. The accumulation of MC3 at the lipid/water interface (Ermilova-Swenson) leads to a reduction of the bilayer thickness not evident from the experiments under present conditions. Surprisingly, the slight and stronger accumulation of MC3 in the center of the bilayer predicted with the Park-Im or the current parameters reproduces the reflectivity profiles equally well, likely due to compensating contributions of MC3 and DOPC. Here, additional high resolution experiments would be valuable to to determine the magnitude of segregation and change in bilayer thickness. In summary, the force field parameters for cationic and neutral MC3 developed in this work provide an accurate model for the ionizable Dlin-MC3DMA lipid in DOPC bilayers and can be used for biomolecular simulations in combination with the AMBER force field for lipids and biomolecules such as proteins, DNA and RNA.

## Author Contributions

N.S. and T.N. designed the research. M.I. and M.H. performed the parametrization of MC3. M.I. performed the simulations and analyzed the data. J.G. and T.N. planned and performed the experiments. N.S., M.I., and J.G. wrote the manuscript. All authors discussed the results and provided feedback on the manuscript.

## Conflicts of interest

“There are no conflicts to declare”.

## Acknowledgements

This research was supported by the Bundesministerium für Bildung und Forschung (BMBF) within the Röntgen-Ångström cluster project Medisoft, number 05K18EZA. The work was also supported by the Emmy Noether program of the Deutsche Forschungsgemeinschaft (DFG, German Research Foundation), number 315221747. We acknowledge the GOETHE HLR for super computing access. M.H. acknowledges support by the German Research Foundation (ProxiDrugs Cluster4Future) and the Max Planck Society. M.I. acknowledges Jürgen Köfinger for fruitful discussions. N.S. thanks Gerhard Hummer for support. M.H thanks Gerhard Hummer for useful discussions. We thank the ISIS Neutron and Muon source (UK) for allocating beamtime at POLREF and Dr Maximilian Skoda for skillful help with the experiments and valuable discussions. The data can be found on https://doi.org/10.5286/ISIS.E.RB2010562

## Notes

### Competing Interest Statement

The authors have declared no competing interest.

### Summary of Updates

Methods from experiments and simulations are updated in the main text. Discussion slightly modified and new figures and tables are added to the supplementary

