## Supplementary Information for "Structural Insights on Ionizable Dlin-MC3-DMA Lipids in DOPC Layers by Combining Accurate Atomistic Force Fields, Molecular Dynamics Simulations and Neutron Reflectivity"

#### 1 Introduction

We derived force field parameters for cationic and neutral MC3 and compared to reflectivity experiments at mixed DOPC/MC3 bilayers. In this supporting information we provide additional information on the methods used in our work.

### 2 Simulation Methods

#### 2.1 Parameterization of MC3 and MC3H

##### 2.1.1 Partial charges

The partial charges in the two protonation states of D-Lin-MC3-DMA were derived following the AMBER lipid14/11 [1,2] charge derivation protocol. In this protocol, the lipid molecule is split into head and tail groups. The groups are capped such that the chemical environment at the split position is preserved during the charge derivation. The charges for the capping groups are constrained to pre-derived values during the charge derivation. The capping molecule and the capped head and tail groups are shown in Figure S1B-D. The partial charges for neutral and cationic MC3 for the different force fields are shown in Figure S2. 20 random conformations were used to obtain the partial charges for the capping group. 100 random capped tail group conformations and 50 random capped head group conformations were used for each protonation state. To obtain the random conformations, 100 ns simulations of a single MC3 or MC3H molecule in water were done using the recently released Park-Im force field parameters [3]. The random conformations were selected from the last 50 ns of the simulations. The geometry of these conformations were further optimized using the Gaussian09 software [4] at the HF/6-31G* level. The RESP [5] charge derivation procedure was used to derive the partial charges for all conformations simultaneously using the RED-vIII.52.pl software [6]. The calculation of the electrostatic potential was evaluated at HF/6-31G*level of theory.

##### 2.1.2 Atom types, angles, dihedrals

In the next step, we assigned the non-bonded parameters namely, atom types, bonds, angles and dihedrals. To that end, a dummy parameter `.itp` file was created for the whole molecule using the `acpype.py` [7] script and assigning atom types according to the GAFF force field [8]. The `.itp` parameter file was then modified by including the correct partial charges (as derived above) and the atom types according to the AMBER Lipid17 force field [9]. The parameter file was then used as input for a custom python script based on MDAnalysis [10] and ParmEd [11] python packages. For a given bond, angle, or dihedral angle, the script assigns the correct values from the AMBER Lipid17 force field (`Lipid17.dat`). If a parameter was not found in `Lipid17.dat`, the parameters from the GAFF2 parameter file `gaff2.dat` were used and the atom types were adjusted accordingly.

#### 2.2 MD simulations

To simulate the MC3-DOPC systems using the Park-Im MC3 parameters and the CHARMM36 force field[12] the initial bilayer configurations were created using the CHARMM-Gui web-server[13]. For simulations with the current MC3 parameters and AMBER Lipid 17 DOPC or the Ermilova-Swenson MC3 and Slipids DOPC, the initial bilayer configurations were created using the the MemGen web-server[14] tool. The systems were solvated with mTIP (CHARMM) or TIP3P water (AMBER, Slipids) with around 72 water/lipid[15,16]. Sodium and chloride ions were added to neutralize the system and to obtain a bulk concentration of 150 mM. The ions were described using recently developed force fields optimized for TIP3P water[17]. The structures were minimized using a gradient descent algorithm followed by 500 ps in the NVT ensemble with an additional flat bottom potential acting on the water molecules to prevent them from penetrating the membrane. The temperature was fixed at 300 K using the velocity rescaling algorithm with stochastic term[18] and with a coupling time constant of 1.0 ps. The system was further equilibrated for 20.0 ns with the semi-isotropic Berendsen barostat[19] and with time constant 5.0 ps. The pressure was fixed at 1.0 bar and the temperature at 300 K using the Berendsen thermostat[19] with time constant 1.0 ps. The production run was carried out for 600.0 ns. The pressure was fixed at 1.0 bar using the semi-isotropic Parinello-Rahman barostat with coupling constant 1.0 ps. The temperature was fixed at 300.0 K using velocity rescaling method with stochastic term[18] and with coupling constant 1.0 ps. Van-der-waals interactions were cut-off at 1.2 nm and a force-switch between 1.0 nm and 1.2 nm was used. Electrostatic interactions were evaluated using the Particle Mesh Ewald[20] method and a cutoff of 1.2 nm. All bonds involving hydrogens were constrained using the LINCS[21] algorithm with LINCS order 4. A time step of 2.0 fs was used to integrate the equations of motion. Simulation trajectories were visualized using VMD[22]. All the simulations were performed using GROMACS (version-2018)[23].

In total, we used 5 different force field combinations: (i) Park-Im MC3/MC3H and CHARMM36 DOPC[3], (ii) Ermilova-Swenson MC3[24] and Slipids DOPC[25], (iii) current MC3/MC3H and AMBER Lipid 17 DOPC[9]. The details on the different simulation systems are shown in Table S1.

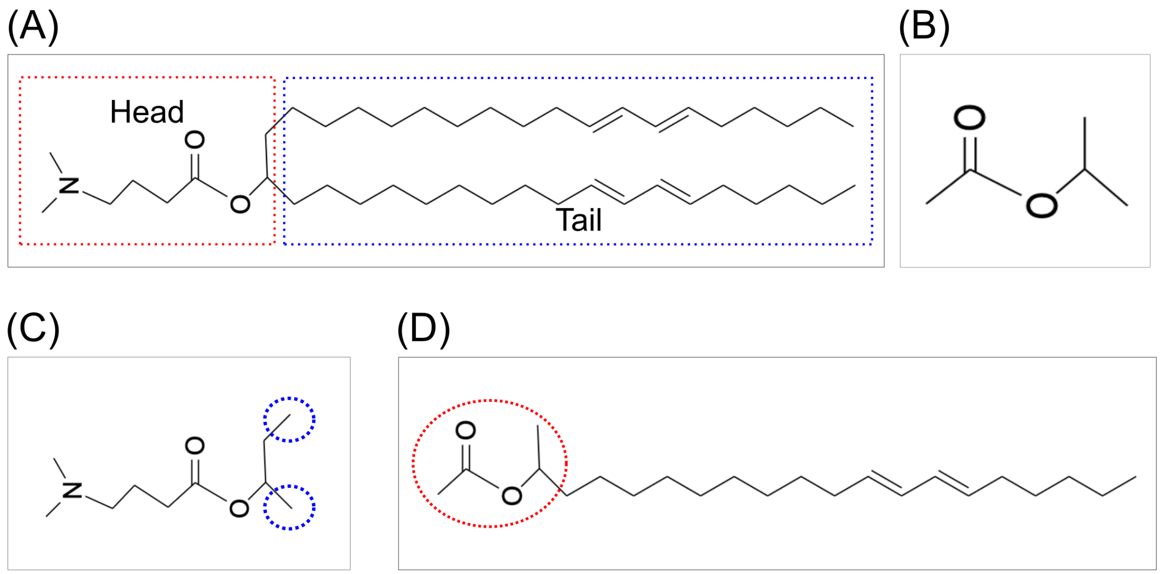


Figure S1: (A) Following the AMBER protocol, the MC3 lipid is divided into a head and tail group. The head and tails are capped during the charge derivation with the capping molecules shown in (B). (C,D) Capped head and tail groups.

| System | MC3/H [mol %] | $N_{DOPC}$ | $N_{MC3}$ | $N_{MC3H}$ | water/lipid | $Na^+$ | $Cl^-$ |
|---|---|---|---|---|---|---|---|
| Pure DOPC† | 0.0% | 200 | 0 | 0 | ∼ 50 | 27 | 27 |
| MC3/DOPC† | 15.0% | 170 | 30 | 0 | ∼ 72 | 40 | 40 |
| MC3H/DOPC† | 15.0% | 170 | 0 | 30 | ∼ 71 | 38 | 68 |
| MC3H/DOPC | 10.0% | 360 | 0 | 40 | ∼ 60 | 105 | 65 |
| MC3H/DOPC | 5.0% | 380 | 0 | 20 | ∼ 60 | 85 | 65 |
| MC3/DOPC | 5.0% | 380 | 20 | 0 | ∼ 60 | 65 | 65 |

Table S1: Simulation setups used in the current work. Systems† were simulated with the three different force fields for neutral MC3 and the corresponding DOPC force field or with the two different force fields for cationic MC3. Here, the box size was about 8.3 nm × 8.3 nm × 10 nm. For all other systems only the current parameters for neutral and cationic MC3 and AMBER Lipid 17 for DOPC were used. Here, the box size was about 11.8 nm × 11.8 nm × 8.7 nm due to the smaller number of ionizable MC3 lipids.

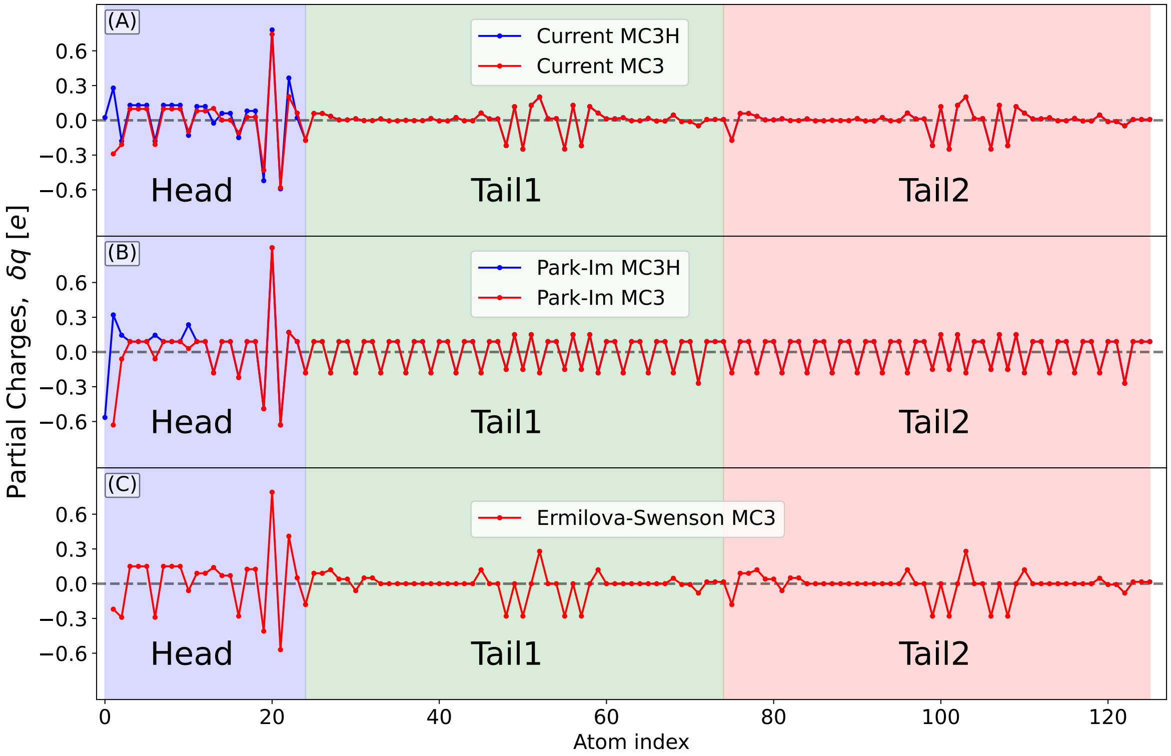


Figure S2: Partial charges for neutral and cationic MC3 from the different force fields. Atom index 0 corresponds to the nitrogen atom in the head group for cationic MC3. Atom index 1 corresponds to the nitrogen atom in the head group for neutral MC3.

#### 2.3 Neutron reflectivity profile from MD simulations

To provide a direct quantitative comparison between experimental and simulation results, the influence of the solid support, which is not present in the simulations, was considered. This is achieved in a three-step procedure: The silicon substrate is modeled by fitting the experimental data of the substrate in solution to a three-slab model (Figure S3). Subsequently, the position of the solid/bilayer interface is determined from the simulations (Figure S4). Finally, we follow the procedure of previous work[26,27] and introduce two physically motivated fit parameters corresponding to the fraction of water in the bilayer leaflet closer to the substrate ($\alpha$) and to the substrate area covered by water patches ($\gamma$).

##### 2.3.1 Modeling of the silicon substrate

The silicon substrate is not present in the MD simulations and is added subsequently to the neutron scattering length density (SLD) by fitting the experimental data of the substrate in solution to a three-slab model. For each experimental sample cell, reflectivity experiments were performed for substrate-$D_2O$ system. The reflectivity profiles were fitted with a three-slab model: Si-layer, $SiO_2$ and $D_2O$. The Refnx[28] python package was used to obtain the substrate SLD profiles. The resulting fits and SLD profiles are shown in Figure S3. Finally, the SLD profiles of the three-slab model were added to the SLD of the simulated free bilayer.

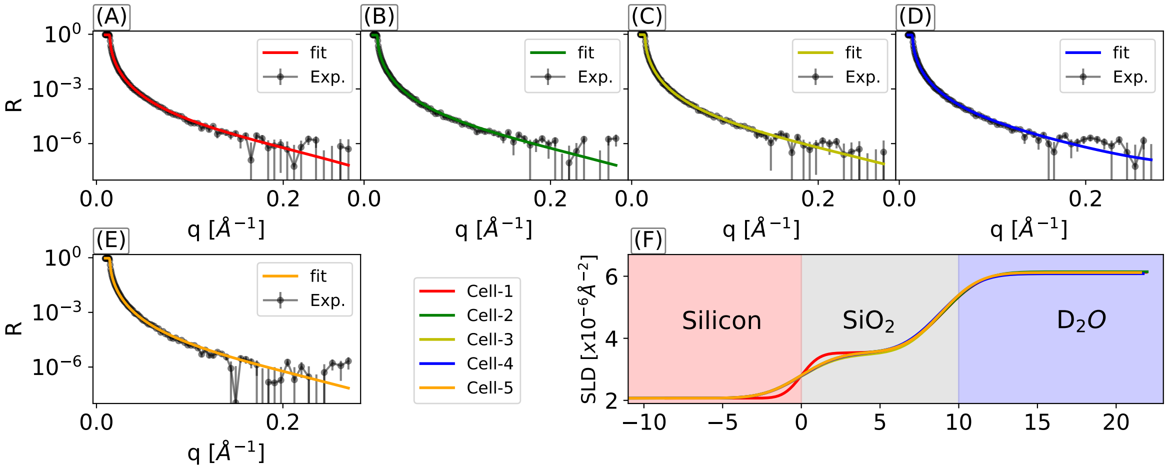


Figure S3: Modeling of the substrate: (A-E) Reflectivity profile $R$ for the silicon/silicon dioxide/$D_2O$ system for each experimental sample cell. (F) SLD corresponding to the best fit of the experimental reflectivity profiles. The substrate thickness in all cases is around 10 Å

##### 2.3.2 Position of the substrate/bilayer interface

The position of the substrate/bilayer interface is determined using the SLD from the free bilayer simulations. Here, the point at which the SLD assumes zero is taken to be the position of the substrate. These points are depicted by vertical dashed lines in Figure S4.

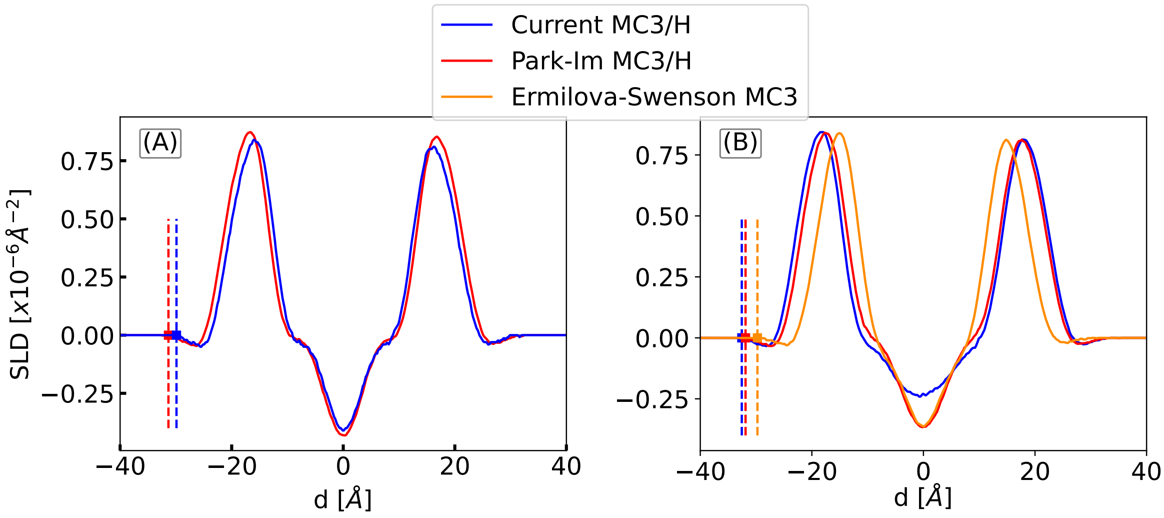


Figure S4: Position of the substrate/bilayer interface. SLD for cationic MC3H/DOPC (A) and neutral MC3/DOPC systems (B) for the different force fields. The vertical dashed lines indicate the position of the substrate, the corresponding distance values are shown in Table S2

| System | Current FF | Park-Im | Ermilova-Swenson |
|---|---|---|---|
| MC3H/DOPC | 29.9 Å | 31.3 Å | N/A |
| MC3/DOPC | 32.5 Å | 31.8 Å | 29.7 Å |

Table S2: Distance of the bilayer/substrate interface from the center of the bilayer for the different force fields. These distances correspond to the position of the vertical dashed lines in Figure S4

.

##### 2.3.3 Fitting of water fraction and water patches

Following previous work[26,27], we introduce two fit parameters. The first parameter, $\alpha$ corresponds to the amount of water in the bilayer leaflet closer to the substrate. It scales only the

SLD of solvent in that leaflet and was chosen to vary between 0 and 1. $\alpha = 1$ corresponds to the unscaled and fully hydrated profile. $\alpha = 0$ corresponds to the fully dehydrated profile (Figure S5B). The second parameter, $\gamma$ corresponds to the fraction of water patches on the substrate. $\gamma = 0$, corresponds to a situation where the substrate is perfectly covered by the bilayer and hence coverage is 100% (Figure S5A).

The total SLD $s(d)$ as a function of the distance $d$ from the center of the bilayer can be written as

$$s(d) = \begin{cases} (1-\gamma)\rho_b(d) + (1-\gamma)\alpha\rho_s(d) + \gamma\rho_{s0} & \text{for} \quad d < d_{\min} \\ (1-\gamma)\rho_b(d) + (1-\gamma)\rho_s(d) + \gamma\rho_{s0}, & \text{otherwise} \end{cases}$$

$\rho_s$ and $\rho_b$ are the SLD of solvent and bilayer, respectively. $\rho_{s0}$ is the SLD of bulk solvent obtained from the fits in Figure S3: $\rho_{s0} = -0.56 \times 10^{-6}$ Å$^{-2}$ for $H_2O$, $\rho_{s0} \sim 6.1 \times 10^{-6}$ Å$^{-2}$ for $D_2O$, and $\rho_{s0} = 2.07 \times 10^{-6}$ Å$^{-2}$ for contrast matched conditions. $d_{\min}$ denotes the position of the minimum in the SLD of the free bilayer (Figure S4). Note that in the center of the bilayer water content is zero over a distance of about 2 nm. Therefore the choice of $d_{\min}$ does not affect the calculation as long as it is in the water-free region of the bilayer.

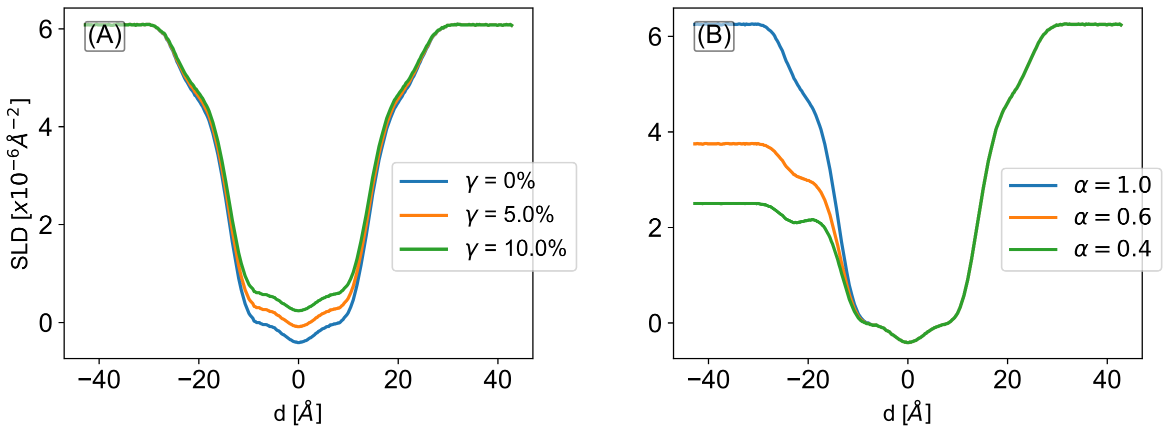


Figure S5: (A) Effect of fit parameter $\gamma$ on the SLD profile. $\gamma = 0\%$ corresponds to the profile obtained from the simulations. (B) Effect of fit parameter $\alpha$ on the SLD profile. $\alpha = 1$ corresponds to the unperturbed SLD profile from the simulations. In both these cases, the solvent is 100% $D_2O$ and the system is the MC3H-DOPC setup with the current force field parameters.

##### 2.3.4 Optimization procedure

To optimize the values of $\alpha$ and $\gamma$ for a given setup, a grid search was used in the parameter space $\alpha \in [0,1]$ and $\gamma \in [0,0.5]$. 10,000 grid points in the ($\alpha$, $\gamma$) plane corresponding to 100 points along each axis were used. At each grid point, we calculated the reduced $\chi^2$ value for the three contrasts

$$\chi^2(\alpha,\gamma) = \frac{1}{(N-1)} \sum_q \frac{(R_q^{\text{exp}} - R_q^{\text{sim}}(\alpha,\gamma))^2}{\sigma_q^2}$$

$R_q^{\text{exp}}$ denotes the reflectivity at point $q$ in the reflectivity profile obtained from the experiments. $R_q^{\text{sim}}(\alpha,\gamma)$ is the reflectivity from the simulations for fixed values of $\alpha$ and $\gamma$. $N$ is the number of data points in the reflectivity profiles and $\sigma_q$ is the error calculated from error propagation of the square root of the number of counts per bin on the detector during the data reduction. The total $\chi^2$ value for fixed values of $\alpha$ and $\gamma$ is the sum over the three different contrasts:

$$\chi^2_{\text{global}}(\alpha,\gamma) = \chi^2_{D_2O}(\alpha,\gamma) + \chi^2_{H_2O}(\alpha,\gamma) + \chi^2_{CMSi}(\alpha,\gamma)$$

The optimal values correspond to the global minimum of $\chi^2_{global}$ on the ($\alpha$, $\gamma$) grid as shown in Figure S6D. The results for the three different force fields are shown in Figure S7.

### 3 Experimental Methods

#### 3.1 Vesicle size for lipid layer deposition

The size of the vesicles with the same lipid compositions used in the lipid layers for both ellipsometry and neutron reflectometry measurements were measured using dynamic light scattering (DLS). Vesicle samples were prepared as per the method described for the inhouse ellipsometry measurements, then were diluted 100x to 0.005 mg/mL lipid in 1x PBS for DLS measurements. The size and polydispersity of the vesicles was measured using a Zetasizer Nano ZS (Malvern Instruments Ltd, UK) with ZEN004 disposable cuvettes filled with 1 mL of diluted vesicles. The data was collected after an equilibration time of 2 minutes at 25 °C and averaged over 3 measurements. The hydrodynamic diameter was calculated using the Einstein-Stokes equation assuming spherical particles (values used for calculation: refractive index of particle = 1.46, refractive index of water = 1.33, viscosity of water = 0.8872 cP).

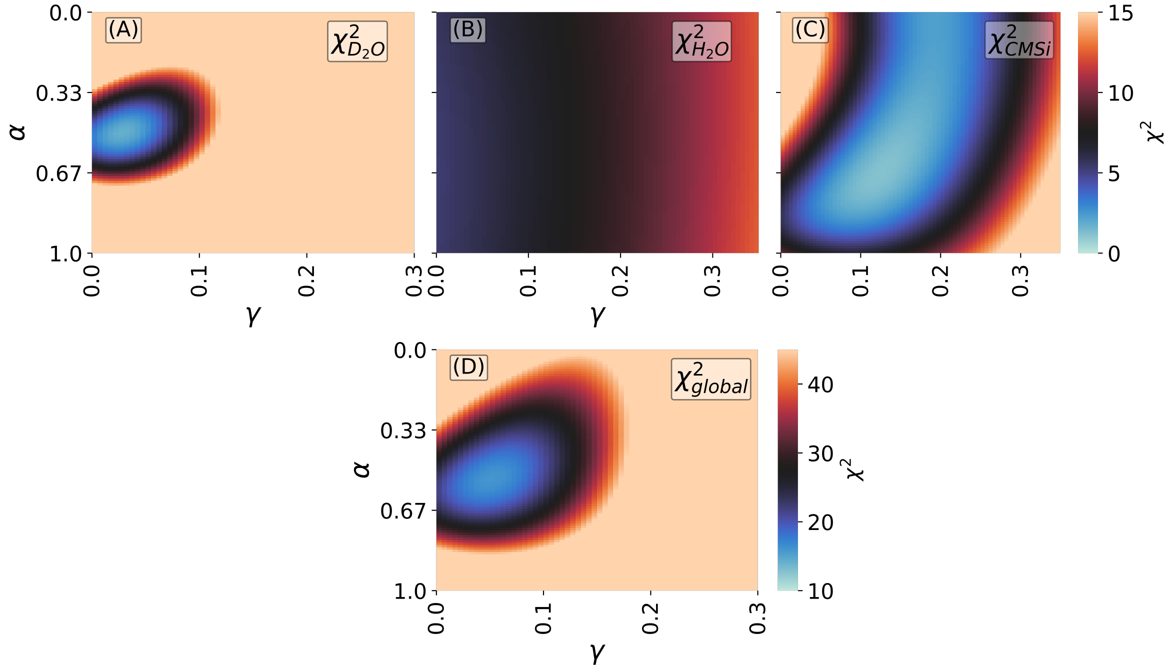


Figure S6: Optimization procedure: At a given point on the ($\alpha$, $\gamma$) grid, $\chi^2$ is calculated for the three contrasts (A-C). The optimal values correspond to the global minimum of $\chi^2_{\text{global}}$ (C). Examples correspond to the MC3H-DOPC setup with the current force field parameters and optimum values are $\alpha = 0.51$ and $\gamma = 0.05$.

| MC3/DOPC | Z-Average (nm) | Number mean (nm) | Polydispersity index (PDI) |
|---|---|---|---|
| 5/95 | 171 ± 2 | 56 ± 15 | 0.47 ± 0.03 |
| 10/95 | 131 ± 1 | 31± 5 | 0.32 ± 0.02 |
| 15/85 | 129± 2 | 38± 3 | 0.26 ± 0.01 |

Table S3: DLS measurements of the vesicles used for deposition of the lipid layer were used to calculate the hydrodynamic diameter. The mean size from the intensity weighted distribution (Z-Average) and number weighted distribution (Number mean) are both presented for clarity.

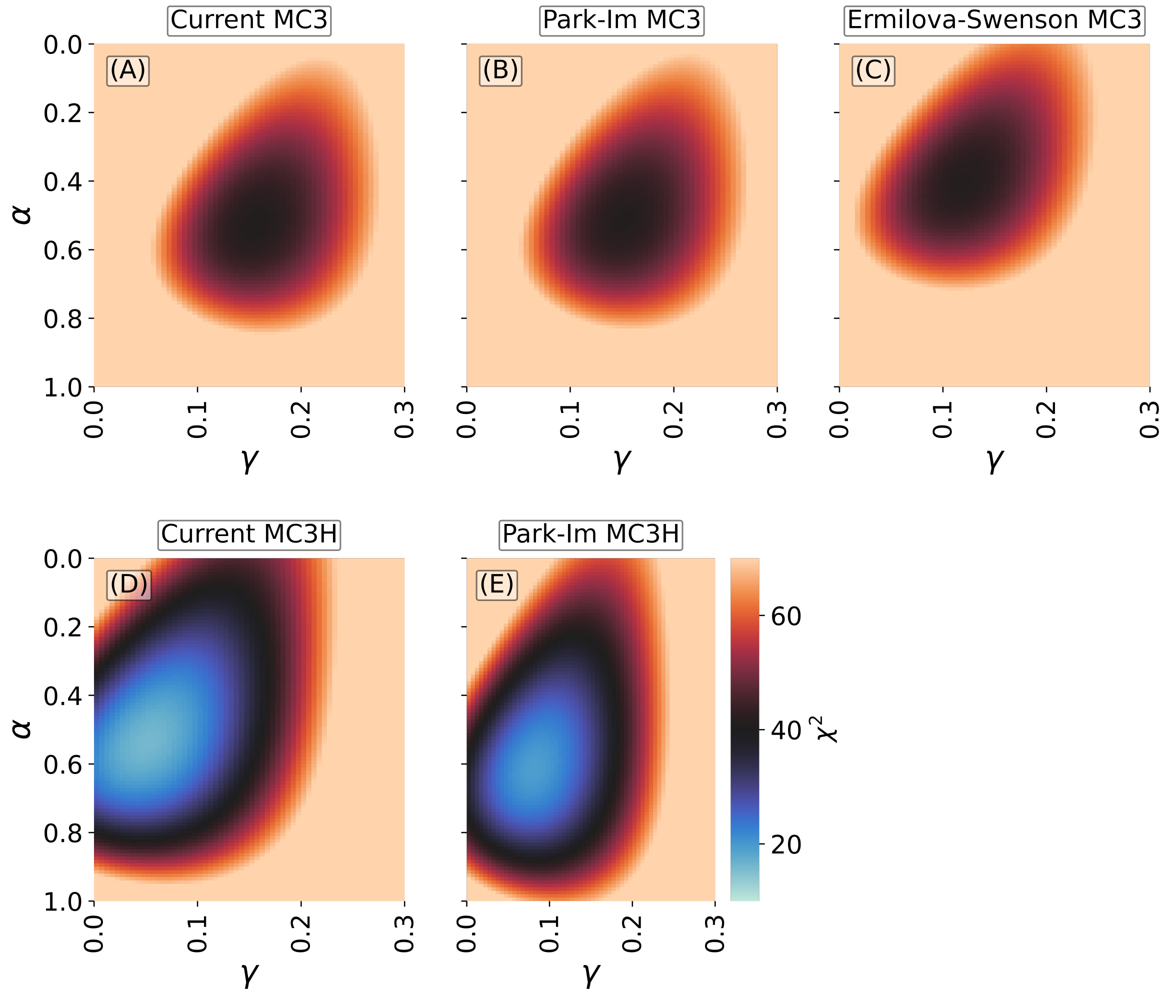


Figure S7: $\chi^2_{global}$ for the different force fields.

### 4 Results

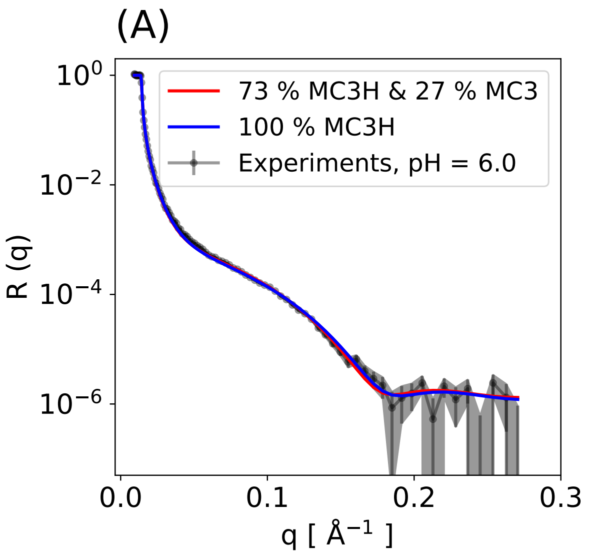


Figure S8: Influence of changes in the protonation degree on the calculated reflectivity profiles. The experimental data are inserted for comparison.

#### 4.1 Simulation convergence

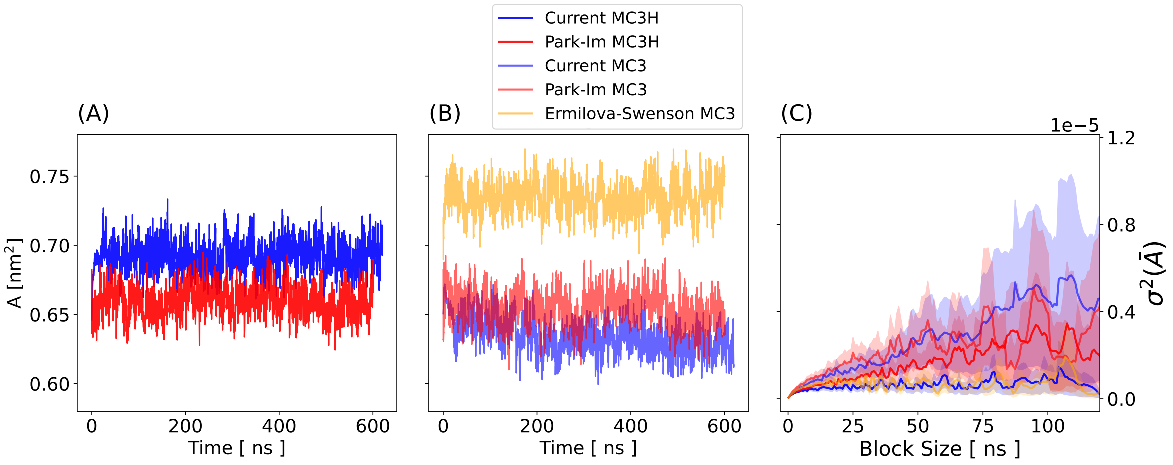


Figure S9: Time trace of the area per lipid,$A$, for the cationic MC3H/DOPC systems (A) and the neutral MC3/DOPC system (B). (C) Result from block averaging for the standard deviation of the ensemble average of the area per lipid ($\sigma^2$) as function of the block size. The shaded region indicates the statistical error of $\sigma^2$.

#### 4.2 Thickness from simulations using different force fields

Here we report characteristic bilayer thicknesses, $D_{HH}$, $2DC$ and $D_B$ (Figure S10). The bilayer thickness, $D_{HH}$ is defined as the distance between the two highest peaks of the total electron density (Figure S10 A). The hydrophobic thickness $2DC$ is defined as the distance between the two points where the DOPC hydrocarbon tails electron density is half of the maximum value[29] (Figure S10 A). The Luzzati thickness ($D_B$), is defined as[25],

$$D_B = d_z - \int_{-d_z/2}^{d_z/2} p(\mathrm{d}) d\mathrm{d} \tag{1}$$

Where $p(d)$ is the water probability distribution along $z$ and is defined as $p(d) = n_w(d)V_w/dV$, where, $n_W$ is the number of water molecules in the slice with volume $dV$, $V_w$ volume of a single water molecule ($V_w \sim 30.5$ Å$^3$). $d_z$ is the z-dimension of the simulation box. Further differences between MC3 added and pure DOPC setups is shown in Figure S11.

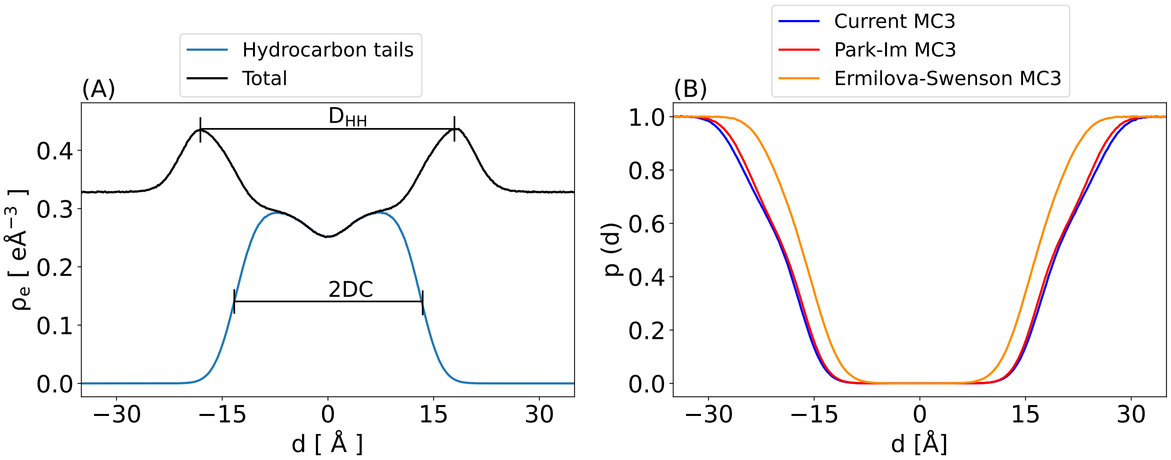


Figure S10: (A) Definition of $D_{HH}$ and $2DC$. (B) Normalized probability distribution of water for the different force fields.

| State | System | Current MC3/H (15%) and Lipid 17 DOPC | Park-Im MC3/H (15%) and CHARMM36 DOPC | Ermilova-Swenson MC3 (15%) and Slipids DOPC |
|---|---|---|---|---|
| Reference | Pure DOPC | 36.8 ±0.6 Å | 38.2±0.6 Å | 36.7±0.3 Å |
| Neutral | MC3/DOPC | 41.5 ±0.2 Å | 39.7±0.8 Å | 33.2±0.4 Å |
| Charged | MC3H/DOPC | 37.4 ±0.6 Å | 38.8±0.8 Å | N/A |

Table S4: Bilayer thickness ($D_{HH}$) for the different systems using different force fields. The bilayer thickness corresponds to the distance between the two highest peaks in the electron density profile. Errors correspond to standard deviations obtained from block averaging (4 blocks for the last 400 ns of the simulations). The reported literature values for pure DOPC from X-ray scattering experiments are 35.3 Å[30], 36.7 Å[31], 36.9 Å[32], 37.1 Å[33] at $T = 30$ °C.

| State | System | Pure DOPC (reference) | 5% MC3/H | 10% MC3/H | 15% MC3/H |
|---|---|---|---|---|---|
| Neutral | MC3/DOPC | 36.8 ±0.6 Å | 37.8±0.3 Å | N/A | 41.5 ±0.2 Å |
| Charged | MC3H/DOPC | 36.8 ±0.6 Å | 36.7±0.4 Å | 36.5±0.4 Å | 37.4 ±0.6 Å |

Table S5: Thickness of DOPC bilayers ($D_{HH}$) containing different mole fractions of cationic or neutral MC3. The bilayer thickness corresponds to the distance between the two highest peaks in the electron density profile. Errors correspond to standard deviations obtained from block averaging (4 blocks for the last 400 ns of the simulations). All simulations were done with the current force fields for cationic and neutral MC3 and the Amber Lipid 17 force field for DOPC.

| State | System | Current MC3/H (15%) and Lipid 17 DOPC | Park-Im MC3/H (15%) and CHARMM36 DOPC | Ermilova-Swenson MC3 (15%) and Slipids DOPC |
|---|---|---|---|---|
| Reference | Pure DOPC | 34.9 ±0.4 Å | 37.9±0.5 Å | 37.3±0.7 Å |
| Neutral | MC3/DOPC | 39.7 ±0.3 Å | 38.7±0.7 Å | 34.1±0.5 Å |
| Charged | MC3H/DOPC | 35.7 ±0.5 Å | 38.6±0.5 Å | N/A |

Table S6: Luzzati thickness ($D_B$) for the different systems using different force fields. Errors correspond to standard deviations obtained from block averaging (4 blocks for the last 400 ns of the simulations).

| State | System | Current MC3/H (15%) and Lipid 17 DOPC | Park-Im MC3/H (15%) and CHARMM36 DOPC | Ermilova-Swenson MC3 (15%) and Slipids DOPC |
|---|---|---|---|---|
| Reference | Pure DOPC | 27.0 ±0.3 Å | 28.9±0.5 Å | 27.9±0.1 Å |
| Neutral | MC3/DOPC | 32.5 ±0.3 Å | 30.6±0.3 Å | 24.3±0.3 Å |
| Charged | MC3H/DOPC | 28.6 ±0.4 Å | 29.9±0.3 Å | N/A |

Table S7: DOPC hydrocarbon thickness (2DC) for the different systems using different force fields. Errors correspond to standard deviations obtained from block averaging (4 blocks for the last 400 ns of the simulations).

#### 4.3 Mass density profiles

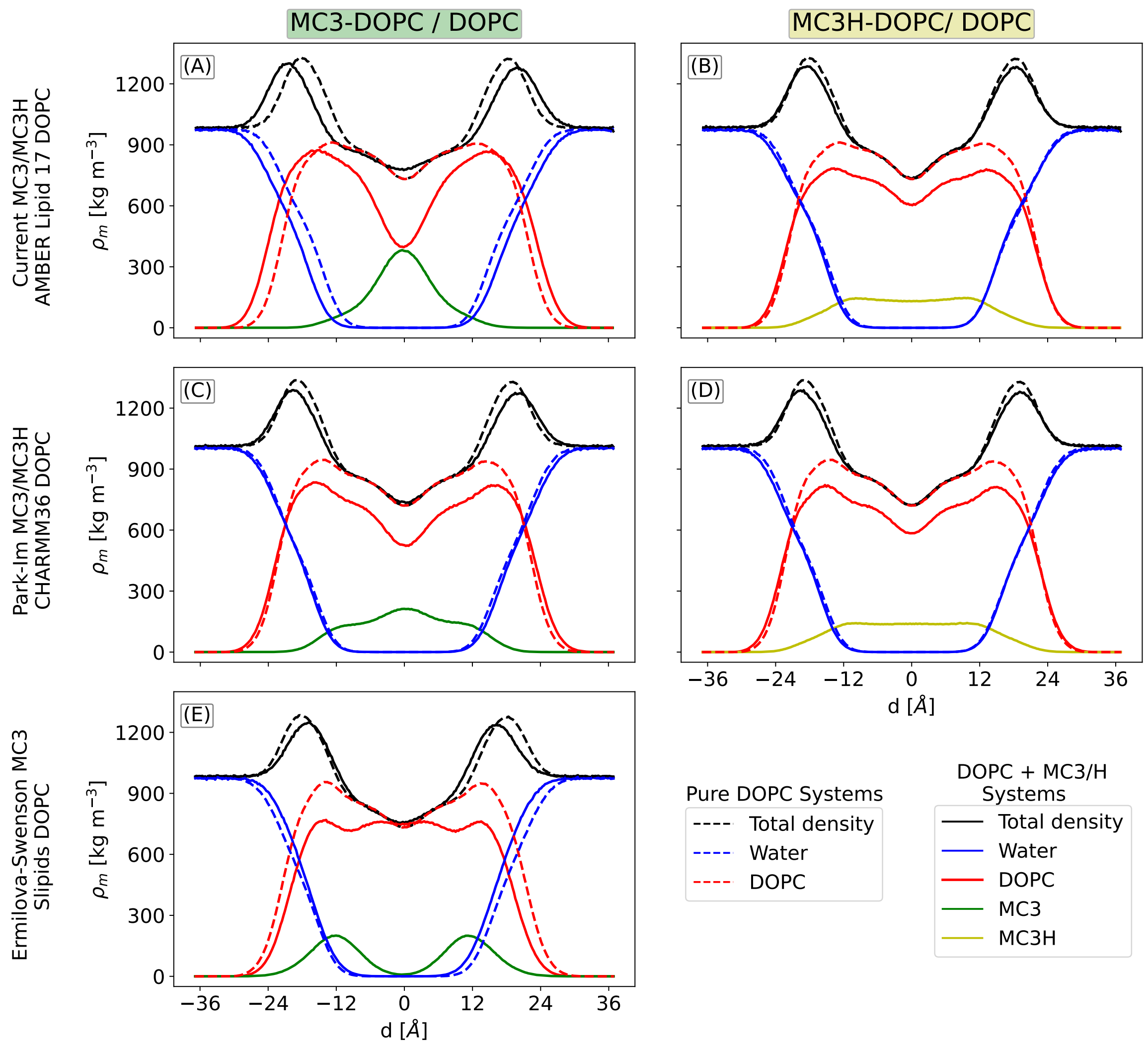


Figure S11: Mass density profiles for pure DOPC systems and systems with mixtures of DOPC and cationic or neutral MC3 for the different force fields. The dashed lines in all cases correspond to a pure DOPC setup and the solid lines correspond to the DOPC-MC3/H setups with 15% MC3/H. The density of neutral MC3 is shown in green, cationic MC3H is shown in yellow, water is shown in blue, and DOPC is shown in red. In each panel, for a given component, the density in both MC3/H-DOPC and pure DOPC systems is plotted.

#### 4.4 Radial distribution functions

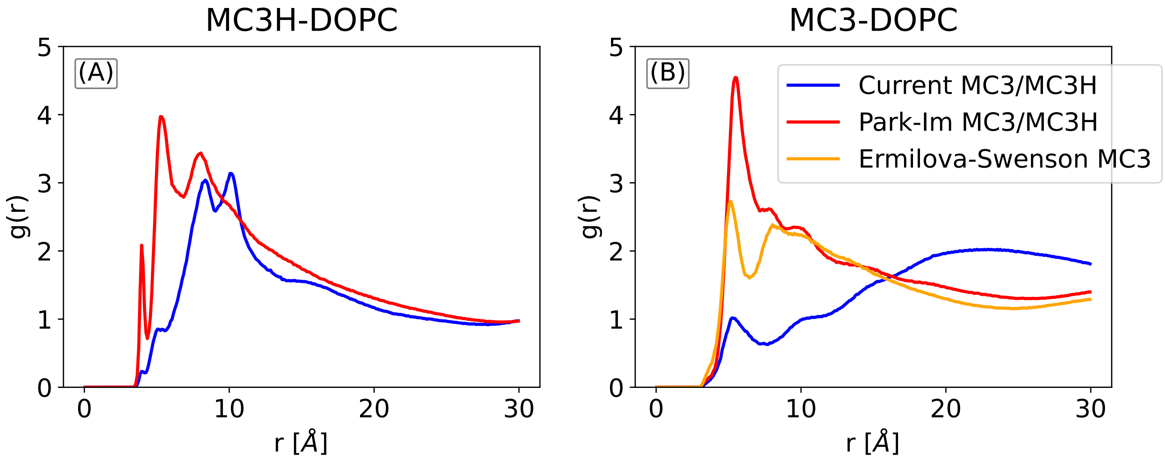


Figure S12: Radial distribution function between the positively charged carbonyl carbon of DOPC and the nitrogen atom of cationic MC3H (A) and neutral MC3 (B) for all force field.

#### 4.5 Experimental reflectivity profiles at pH 6 and pH 7

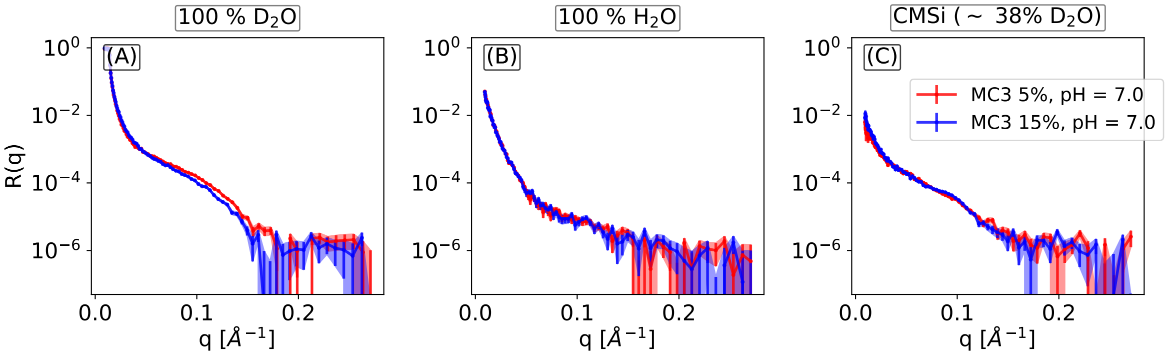


Figure S13: Comparison of the experimental reflectivity profile at pH 6 and pH 7 for the DOPC bilayer with 15% MC3. The shift of the characteristic minimum is clearly visible at 100% $D_2O$ and indicates an increase of the bilayer thickness for uncharged MC3 at pH 7 in agreement with the simulations (Table S5).

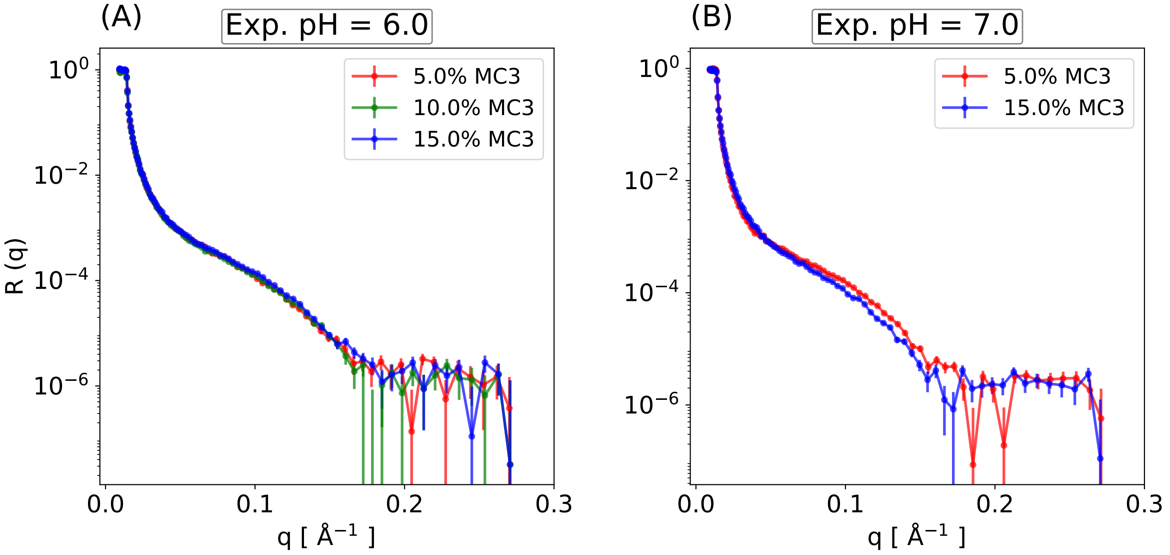


Figure S14: Comparison of the experimental reflectivity profile at different MC3 fractions for pH 6 and pH 7. The profiles correspond to 100% $D_2O$ solvent contrast. For pH 7, the shift of the characteristic minimum to lower $q$ upon increasing the MC3 fraction is clearly visible.

#### 4.6 Reflectivity profiles at different MC3 fractions

##### 4.6.1 5% MC3H-95% DOPC

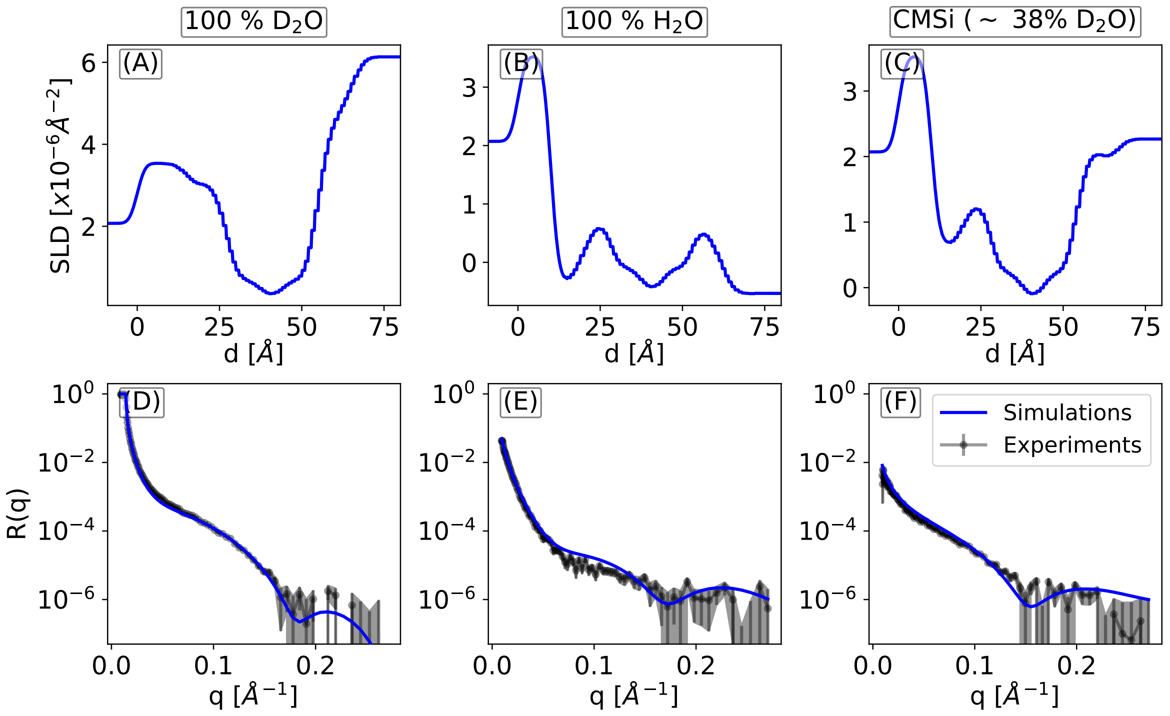


Figure S15: Scattering length density profiles from simulations at different deuteration levels (top) and corresponding reflectivity profiles (bottom). The systems contain 5% MC3H and 95 % DOPC. Global fitting to the three contrasts yields $\alpha = 0.51$ and $\gamma = 11.6\%$. The simulations were done with the current force fields for MC3 and the AMBER Lipid 17 force field for DOPC. The experiments were performed at pH=6.0.

###### 4.6.2 10% MC3H-90% DOPC

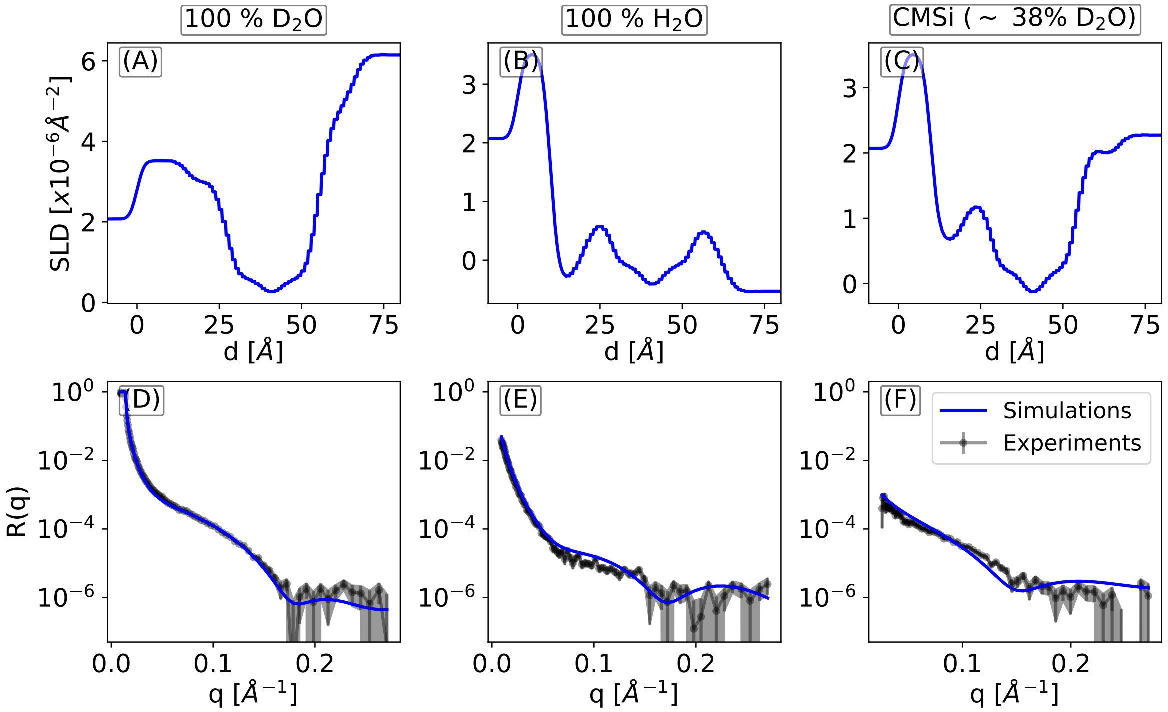


Figure S16: Scattering length density profiles from simulations at different deuteration levels (top) and corresponding reflectivity profiles (bottom). The systems contain 10% MC3H and 90 % DOPC. Global fitting to the three contrasts yields $\alpha$ = 0.52 and $\gamma = 10.1\%$. The simulations were done with the current force fields for MC3H and the AMBER Lipid 17 force field for DOPC. The experiments were performed at pH=6.0.

##### 4.6.3 5% MC3-95% DOPC

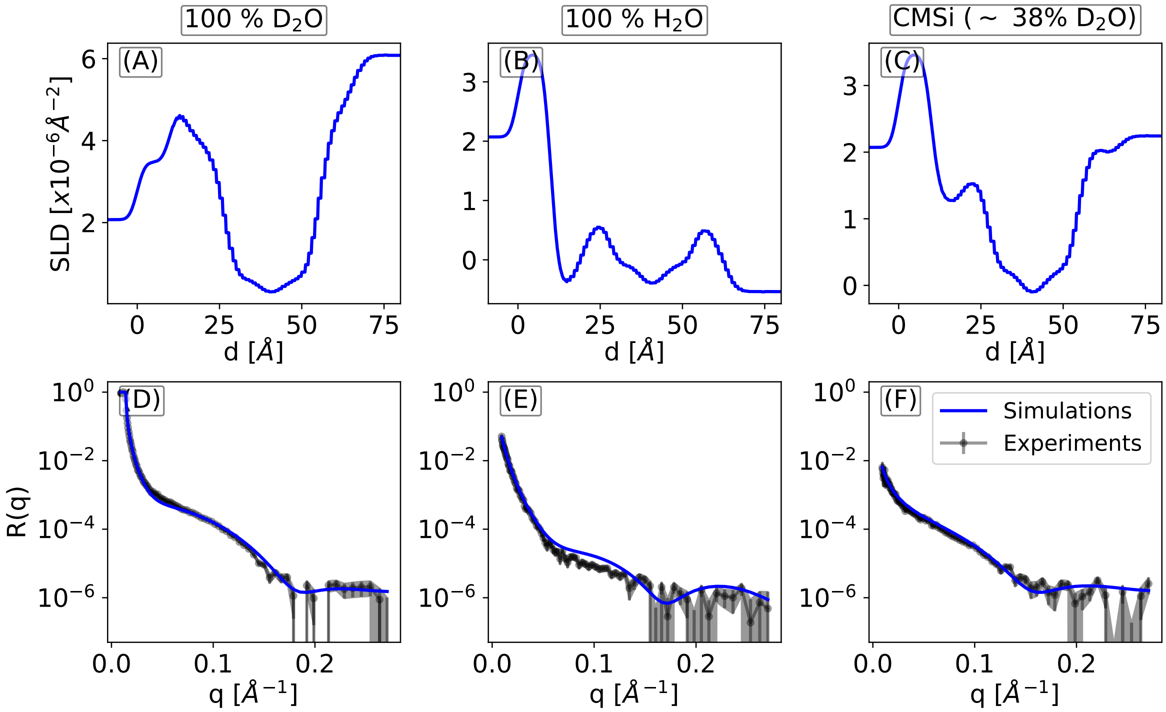


Figure S17: Scattering length density profiles from simulations at different deuteration levels (top) and corresponding reflectivity profiles (bottom). The systems contain 5% MC3 and 95 % DOPC. Global fitting to the three contrasts yields $\alpha = 0.77$ and $\gamma = 10.6\%$. The simulations were done with the current force fields for MC3 and the AMBER Lipid 17 force field for DOPC. The experiments were performed at pH=7.0.

#### 4.7 Form factors from the MD simulations

The form factors were calculated from the Fourier transform of the electron density profiles obtained from the simulations via

$$F(q) = \int_{-\infty}^{+\infty} (\rho_e(z) - \rho_b) e^{-iqz} dz \tag{2}$$

$\rho_e$ is the electron density along bilayer normal. $\rho_b$ is the bulk electron density. The integral was evaluated using standard integral tools available in the `Numpy` python package.

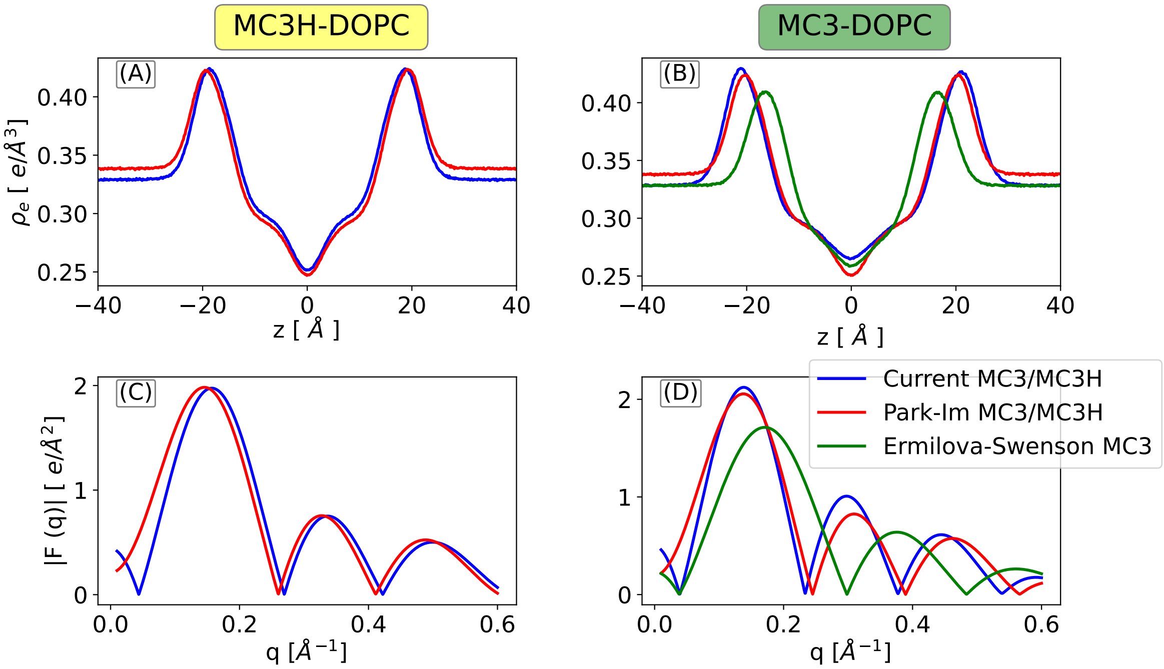


Figure S18: X-ray form factors from the simulations with the different force fields. (A, B) Electron densities for MC3H-DOPC and MC3-DOPC systems. (C, D) Calculated X-ray scattering form factors $F(q)$.
